# Mapping human natural killer cell development in tonsil

**DOI:** 10.64898/2026.05.13.722762

**Authors:** Everardo Hegewisch-Solloa, Janine E. Melsen, Ansel P. Nalin, Hiranmayi Ravichandran, André F. Rendeiro, Bethany Mundy-Bosse, Johannes C. Melms, Shira E. Eisman, Benjamin Izar, Eli Grunstein, Thomas J. Connors, Olivier Elemento, Aharon G. Freud, Amir Horowitz, Emily M. Mace

**Affiliations:** Department of Pediatrics, Vagelos College of Physicians and Surgeons, Columbia University Irving Medical Center, New York NY 10032; Department of Immunology, Leiden University Medical Center, Leiden, The Netherlands; Laboratory for Pediatric Immunology, Willem-Alexander Children’s Hospital, Leiden University Medical Center, Leiden, The Netherlands; Department of Pathology, The Ohio State University, Columbus, OH 43210, USA; Comprehensive Cancer Center and The James Cancer Hospital and Solove Research Institute, The Ohio State University, Columbus, OH 43210; Department of Physiology, Biophysics and Systems Biology, Weill Cornell Medicine, New York, NY, 10065; Caryl and Israel Englander Institute for Precision Medicine, Weill Cornell Medicine, New York, NY, USA; CeMM Research Center for Molecular Medicine of the Austrian Academy of Sciences, Lazarettgasse 14 AKH BT 25.3, 1090, Vienna, Austria; Ludwig Boltzmann Institute for Network Medicine at the University of Vienna, Augasse 2-6, Vienna, A-1090, Austria; Division of Hematology, Department of Internal Medicine, The Ohio State University, Columbus, OH 43210, USA; Comprehensive Cancer Center and The James Cancer Hospital and Solove Research Institute, The Ohio State University, Columbus, OH 43210; Department of Medicine, Division of Hematology/Oncology, Columbia University Irving Medical Center, New York, NY, 10032; Columbia Center for Translational Immunology, Columbia University Irving Medical Center, New York, NY, 10032; Herbert Irving Comprehensive Cancer Center, Columbia University Irving Medical Center, New York, NY, 10032; Program for Mathematical Genomics, Columbia University, New York, NY, 10032; Department of Otolaryngology - Head and Neck Surgery, Columbia University Medical Center, New York, New York 10032; Department of Pediatrics, Division of Pediatric Critical Care and Hospital Medicine, Columbia University Irving Medical Center, New York, NY 10024; Institute for Computational Biomedicine, Weill Cornell Medicine, New York, NY, 10065; Department of Oncological Sciences, Precision Immunology Institute, Tisch Cancer Institute, Icahn School of Medicine at Mount Sinai, New York, NY, 10029

**Keywords:** natural killer cell, cyclic immunofluorescence

## Abstract

Secondary lymphoid tissue, including tonsil, supports human NK cell development, but the spatial organization and tissue niches that drive this differentiation remain undefined. Here, we used single cell analysis of cyclic immunofluorescence to generate a comprehensive atlas of human NK cell development in tonsil. By integrating regional localization, chemokine signaling, cytokine availability, and cell phenotype, we show that NK cell differentiation follows a reproducible spatial trajectory defined by stage-specific cell-cell interactions. CD34+ NK cell progenitors are found in the interfollicular domain in proximity to high endothelial venules, consistent with this route of progenitor entry into tissue. Mature NK cells are primarily found in the T cell-rich parafollicular domain, where they interact with other NK cells and T cell subsets. Local inflammation increases NK cell frequency in tissue through both proliferation of NK progenitors and increased frequency of mature NK cells. Finally, we identify a subset of tonsil stromal cells that support differentiation of NK cells in vitro and proliferation of NK precursors in situ. Together, these findings demonstrate that spatial localization defines human NK cell development and provide an in situ definition of niches that support human NK cell differentiation in tonsil.

## Introduction

Human secondary lymphoid tissue, including tonsil, generates and coordinates immune responses. This coordination depends on the spatial organization of immune and stromal cells into defined microdomains that enable stage-specific communication and differentiation. While these spatial relationships have been mapped in mouse, comparable studies in human SLT remain scarce and rarely resolve cellular niches that support immune cell differentiation. This gap is particularly limiting for understanding human NK cell development, which occurs in SLT but lacks a stage-resolved spatial framework.

Human natural killer (NK) cell development is defined by six sequential stages characterized by lineage potential, cell surface protein expression, and function (Fig. S1A) (1–6). The isolation of multipotent NK/ILC progenitors that differentiate into mature NK cells or helper ILCs in vitro from secondary lymphoid tissues (SLT) and the presence of NK cell developmental stages in tonsil demonstrates that tonsil serves as a site of NK/ILC development (3, 4, 7, 8). However, whether discrete stages of NK cell development occupy distinct locations within tonsil, what cellular interactions support each stage, and how this organization responds to perturbation remain unresolved.

Prior work has localized CD34+CD45RA+ NK cell progenitors (NKP) and mature NK cells to the parafollicular and subepithelial domains of tonsil, though discrete developmental stages have not been resolved in situ (8–11). The niche elements thought to support NK cell development, including gp38+ fibroblastic reticular cells (FRCs), CD31+ endothelial cells that form high endothelial venules (HEVs), and cytokines including IL-15, IL-7, stem cell factor (SCF), FMS-like tyrosine kinase-3 ligand (Flt3L), and Delta-like canonical notch ligand-1 (DLL1), have been largely characterized from in vitro studies, where direct contact with stromal cells is required to promote NK cell development and proliferation (12–18). How these signals are spatially localized in functional niches that support NK cell development in tissue has not been determined.

Using cyclic immunofluorescence (CyCIF) microscopy, we generated a spatial map of immune cell populations in human tonsil. We used this map to define the in situ trajectory of NK cell development, identify and functionally validate stromal cell niches that support NK cell differentiation, and determine how this organization responds to local inflammation. Together, these findings define the cellular niches that support human NK cell development in tonsil while providing insight into the spatial organization of human secondary lymphoid tissue.

## Results

### Distribution of NK cells and other immune subsets in human tonsil

To map NK cell developmental subsets in human we used CyCIF microscopy on FFPE sections from 10 pediatric donors, capturing up to 43 unique markers across 23 rounds of iterative staining (Fig. 1A) (19, 20). Following segmentation with Mesmer (21), tonsil neighborhoods were assigned to 4 anatomical domains (follicle, T cell zone, epithelial and interfollicular) using CD20, CD3, pan-keratin, and collagen type I (Fig. 1A). Single cell classification using 25 markers identified 6 major immune cell subsets (Fig. 1B), which we further refined to 27 subpopulations (Fig. S1C). NK cell developmental subsets were defined by phenotype, proliferation, and tissue residency: stage 1/2 NK progenitors (NKPs; Lin–CD45+CD34+CD122+CD127+CD49d+), stage 3 (immature NK; Lin–CD45+CD56+CD127+GZMK–GZMB–), stage 4 (Lin–CD45+CD56+CD127+/–GZMK+/–GZMB–), stage 5 (Lin–CD45+CD56+NKp46+CD127–GZMB+CD57–), tissue-resident (tr) stage 4 and 5 NK cells (CD69+ or CD49a+), and proliferating subsets (Ki67+) (Fig. 1D, E; Fig. S1A–D). Lin–CD45+ cells that did not fall into one of the NK stage-specific gates were classified as NKp46+CD56+ NK cells (Fig. 1D, E; Fig. S1A-D). NK cells made up 0.35% to 3.5% of total cells across donors (Fig. 1C), with CD127+ immature NK cells and CD56+NKp46+ NK cells most abundant in 5 donors selected for further analysis (Fig. 1D).

**Fig. 1.**
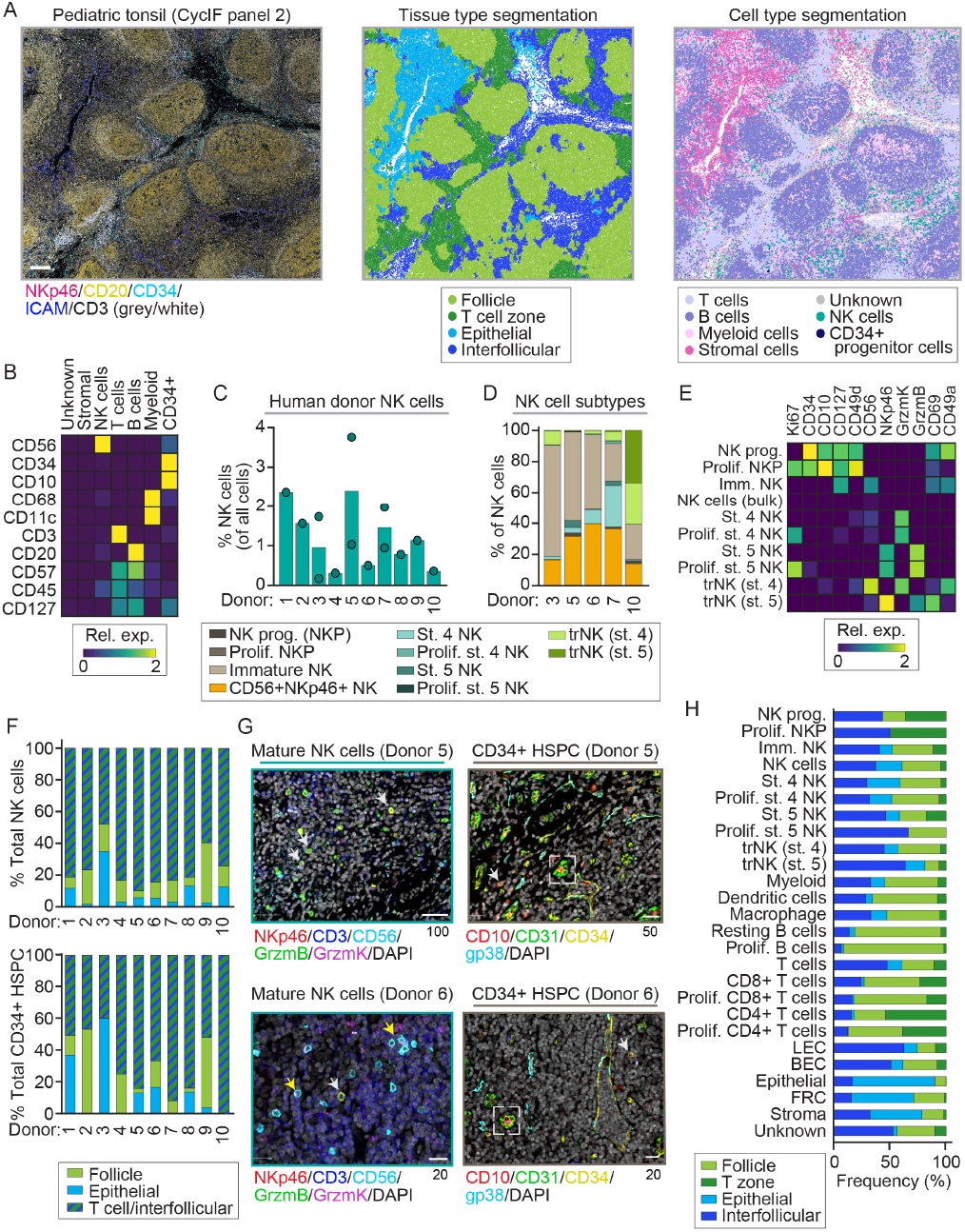
Spatial localization of NK cell subsets in tonsil defined by CyCIF. Representative cyclic immunofluorescence (CyCIF) image (left, CyCIF panel 2 [Supp. Table S1]), automated domain classification (HALO AI, middle), and cell phenotype mask from supervised cell gating (right) of a pediatric tonsil FFPE sample stained for 43 markers over 23 cycles. CD20 (yellow), CD34 (cyan), NKp46 (magenta), ICAM (blue), and CD3 (grey/white); scale bar=100 µm. B) Heatmap of mean lineagespecific marker expression from pediatric tonsil (n=9 donors; 9 ROIs; CyCIF panel 2). C) Percent NK cells of total cells per ROI (n = 10 donors, 12 ROIs; bars represent mean per donor, dots represent individual ROIs). D) Frequency of NK cell subsets within CD34+Lin– and CD56+Lin–compartments (n = 5 donors, 5 ROIs). Subset color legend below: NKP (dark brown), proliferating NKP (Prolif. NKP, brown), immature NK (CD127+, taupe), CD56+NKp46+ NK (orange), stage (St.) 4 NK (aqua), proliferating St. 4 NK (GrzmK+, light teal), St. 5 NK (medium teal), Prolif. St. 5 NK (dark teal), tissue-resident NK stage 4 (trNK st. 4, celery green), and trNK st. 5 (olive green). E) Heatmap of mean marker expression across NK cell developmental subsets (n = 5 donors, 5 ROIs). F) Distribution of NK cells (top) and CD34+ HSPCs (bottom) across tonsil microanatomical domains (n = 10 donors, 12 ROIs; percentages averaged for donors with >1 ROI). G) Representative CyCIF images of mature NK cells (left) and CD34+ HSPCs (right) from two donors. Top row, donor 5; bottom row, donor 6. Left panels: NKp46 (red), CD3 (blue), CD56 (cyan), GrzmB (green), GrzmK (magenta), DAPI (grey/white); white arrowheads, St. 5 NK cells; yellow arrowheads, St. 4 NK cells. Right panels: CD10 (red), CD31 (green), CD34 (yellow), gp38 (cyan), DAPI (grey/white); white arrowheads, CD34+ HSPC; white dashed boxes, high endothelial venules (HEVs). Scale bars: top row = 100 µm (left) and 50 µm (right); bottom row = 20 µm. (H) Distribution of immune cell populations across tonsil microanatomical domains (n = 5 donors, 5 ROIs). All panels show CyCIF data. Two CyCIF cohorts were used: a 5-donor cohort (panels A, D, E, G, H) and a 9-donor cohort (panel B). Panels C and F combine both cohorts (10 unique donors, 12 ROIs)

NK cell developmental subsets occupied distinct anatomical domains within tonsil. Mature stage 4 and stage 5 NK cells localized predominantly to the interfollicular domain and T cell zone (Fig. 1F, G; Fig. S2A, B). CD34+ NKPs were also enriched in the interfollicular domain, where they were observed adjacent to high endothelial venules and lymphatic vessels (Fig. 1G; Fig. S2C, D), consistent with entry into tonsil through these structures. Manual annotation and automated domain classification (HALO AI) of tonsil domains gave concordant results, with the trained classifier (5 donors, 5 ROIs) resolving finer spatial relationships among NK subsets (Fig. 1A, H). Tissue-resident stage 5 NK cells preferentially localized to the interfollicular domain relative to tissue-resident stage 4 NK cells, while non-resident stage 4 and stage 5 NK cells were enriched in the T cell zone, with GrzmB+ stage 5 NK cells the most abundant population there (Fig. 1G, H). Less mature NK cell populations were more broadly distributed across both domains.

### Chemokine signaling positions NK cell developmental subsets in distinct tonsil domains

NK cell developmental subsets expressed stage-specific chemokine receptors that suggested distinct trafficking and localization signals. Analysis of previously published RNA-seq data from sorted stages 3-5 tonsil NK cells (22) showed that early stages (3 and 4A) had higher expression of CXCR6, CCR6, CCR4, and CXCR5, while later stages (4B and 5) had increased CX3CR1, CCR1, CCR5, and CXCR3 (Fig. 2A, Fig. S3). To extend this to earlier developmental stages and to confirm protein expression, we performed flow cytometry on paired peripheral blood (PB) and tonsil samples from 6–7 pediatric donors (Fig. 2B, Fig. S4A–C). Stage 1 NKPs in tonsil expressed CCR6, CXCR4, and CXCR5 at higher levels than mature NK subsets, and CCR5 was uniquely expressed by stage 2 NKPs in tonsil but not in blood. Stages 4B–6 showed increased CXCR3, and CX3CR1 was detected on a subset of stage 6 NK cells in tonsil and on stages 4–6 in blood. Tonsil and blood NK cells showed distinct receptor profiles overall, particularly at CCR4, CCR5, CCR6, CXCR4, CXCR5, and CXCR6 (Fig. 2B, Fig. S4B, C).

**Fig. 2.**
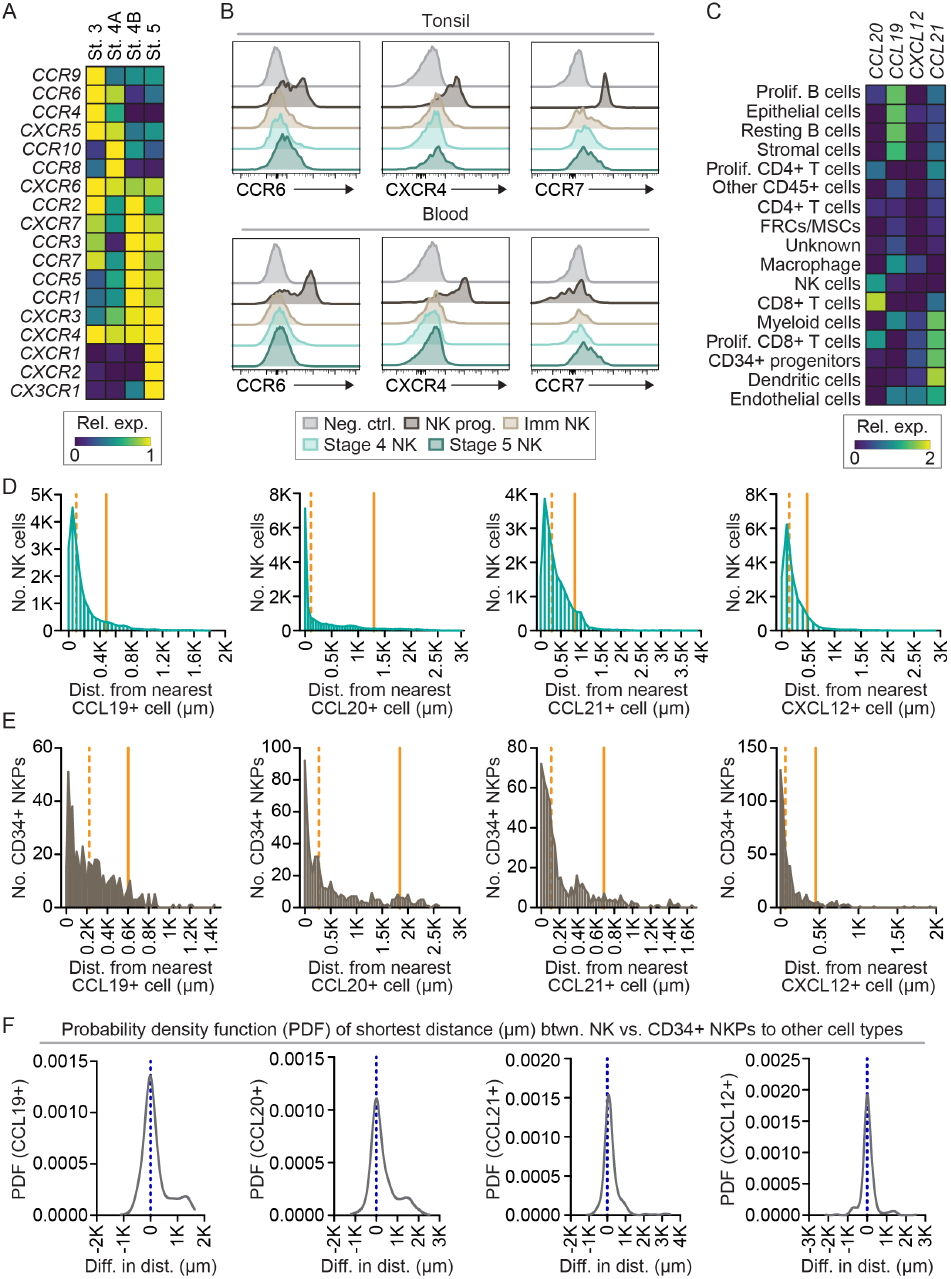
Chemokine receptor expression and localization of NK cells in tonsil. A) Normalized gene expression from bulk RNA-seq of sorted NK cell developmental subsets (22). Heat map=row scaled FPKM. n=12 donors. See also Fig. S1 for developmental stage definitions. B) Chemokine receptor expression on NK cell subsets from tonsil (top) or peripheral blood (bottom). FACS panels listed in Supp. Table S3. Representative of n= 6-7 donors per panel. C) Relative fluorescence intensity (expression) of CCL19, CCL20, CCL21, or CXCL12 from CycIF imaging (n=9 donors; 9 ROIs). D) Median shortest distance (µm) of CD56+LinNK cells to CCL19+, CCL20+, CCL21+, or CXCL12+ cells in tonsil (n=9 donors; 9 ROIs). E) Median shortest distance (µm) of CD34+ progenitor cells to CCL19+, CCL20+, CCL21+, or CXCL12+ cells in tonsil (n=9 donors; 9 ROIs). F) Comparison of differences in shortest distance (µm) of NK cell and CD34+ progenitors to CCL19+, CCL20+, CCL21+, and CXCL12+ cells calculated by probability density function (PDF) from CyCIF data (n=9 donors; 9 ROIs).

We next asked whether spatial localization of NK cell subsets corresponded to the distribution of their chemokine ligands. Fluorescence intensity from CyCIF identified CCL19, CCL20, CCL21, and CXCL12 in distinct tonsil domains across 9 donors (Fig. 2C), with CCL19 enriched on B cells, CCL20 on T cells, and CCL21 and CXCL12 on stromal and endothelial populations. Mature NK cells were closer to CCL19+ and CCL20+ cells than CCL21+ and CXCL12+ (Fig. 2D), consistent with their enrichment in T cell and B cell domains. In contrast, CD34+ NK progenitors were closer to CXCL12+ and CCL21+ (Fig. 2E), consistent with localization near HEVs and stromal cells in the interfollicular domain. To compare the relative shortest distance of mature NK cells and NK progenitors to chemokine positive cells, we calculated the difference in shortest distance of our populations of interest to CCL19-, CCL20-, CCL21-, and CXCL12positive cells. To run this analysis, we filtered our data to include equal numbers of mature NK cells and CD34+ progenitors. Mature NK cells had a higher likelihood of being in proximity to CCL19 and CCL20 than CD34+ NK progenitors (Fig. 2F). This pattern was not observed for CCL21 and CXCL12, where CD34+ cells were slightly more likely to be in closer proximity (Fig. 2F).

### Stage-specific changes in cell-cell interactions of NK cell developmental subsets

Having quantified the spatial localization of NK cell subsets, we focused on their interactions with other immune cells. We mapped interactions between NK cells and other immune subsets. Stage 4 and 5 NK cells were often visualized in proximity to CD4+ and CD8+ T cells in the interfollicular and parafollicular domains (Fig. 3A). Interactions between CD34+ progenitors and other cell types were much rarer due to the low frequency of these cells, but they could be visualized in the T cell zone (interfollicular/parafollicular) (Fig. 3B).

**Fig. 3.**
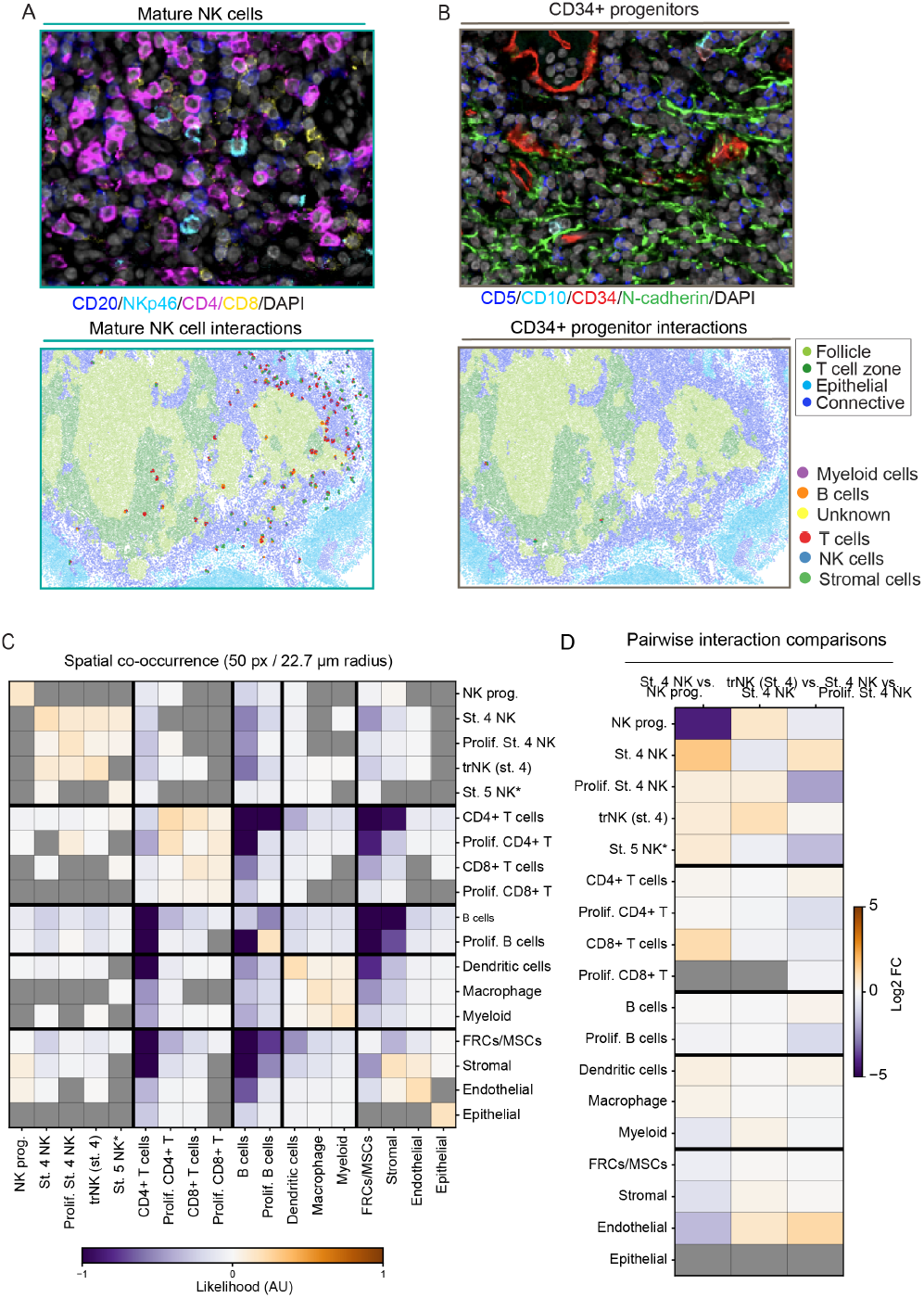
NK cell interactions in tonsil change through development. A)Representative CyCIF image of NK cells interacting with CD3+ T cells (top) and representative map of interactions between NK cells and other cell types (T cells=red, other NK cells=blue, stromal cells=green, myeloid cells=purple, B cells=orange, unknown cells=yellow). Cells were segmented and regions of tonsil were classified by CyCIF staining (light green=follicle, dark green=T cell zone, light blue=epithelial, dark blue=interfollicular). B) Representative CyCIF image of NK cells interacting with CD34+ HSPCs (top) and interaction map (bottom). Representative of 10 ROIs from 9 donors. C) Interaction analysis showing populationlevel likelihood of interaction between subsets (-1.0=negative likelihood, 0=neutral, +1.0=positive likelihood, grey=n.s.; n=9 donors; 9 ROIs). D) Pairwise comparison of the likelihood of interaction between NK cell subsets (Y-axis) and other cell subsets (X-axis) (-6=negative likelihood, 0=neutral, +6 = positive likelihood; n=9 donors; 9 ROIs)

We then performed pairwise radius spatial co-occurrence analysis to identify phenotypes that colocalize more or less frequently within a 50-pixel radius than expected by chance (23). This analysis showed that NK developmental subsets clustered preferentially with one another, with Stage 4 NK cells co-occurring with proliferating Stage 4 NK, tissue resident Stage 4 NK, and the pooled Stage 5 NK population at frequencies higher than expected by chance (Fig. 3C). NK progenitors also showed homotypic clustering and were enriched for proximity to stromal cells. In contrast, NK cell subsets showed negative enrichment for interactions with B cell populations, consistent with their relative exclusion from B cell follicles.

Pairwise comparison between subsets identified reorganization of cell-cell interactions throughout NK cell developmental trajectory (Fig. 3D). The transition from NK progenitor to Stage 4 NK was accompanied by a shift away from stromal and endothelial neighborhoods toward increased homotypic Stage 4 contacts and enrichment for CD8+ T cells, suggesting that lineage commitment includes relocalization out of stromal and endothelial cell neighborhoods. Tissue residency at Stage 4 was associated with modest changes overall, with the strongest signal being enrichment of tr-Stage 4 NK cells near endothelial cells and other tr-Stage 4 cells, indicating that acquisition of tissue residency at this developmental stage is not associated with large changes in cellular localization. Finally, we compared non-proliferating to actively proliferating Stage 4 NK cells and observed a strikingpattern: proliferating Stage 4 NK cells were preferentially located near other proliferating cells, including proliferating B cells, proliferating CD4+ T cells, and NK progenitors, while non-proliferating Stage 4 NK cells were enriched near endothelial cells. This suggests the existence of shared proliferation niches in the tonsil that bring actively dividing lymphocyte subsets into proximity with each other.

Together, these analyses indicate that NK development in tonsil is associated with spatially distinct neighborhoods. The earliest CD34+ progenitors occupy a stromal/endothelial niche; commitment to Stage 4 is accompanied by relocalization into homotypic and CD8+ T cell-rich domains; and active proliferation, regardless of developmental stage, is associated with co-localization in shared proliferative compartments.

### NK progenitors proliferate adjacent to IL15+ cells in tonsil

Our spatial co-occurrence analyses suggested that actively proliferating cells occupy shared niches in the tonsil, raising the question of which developmental stages of NK cells are most proliferative and what cues might support their division in tissue. To address this, we first characterized proliferative subsets during NK cell differentiation using an in vitro co-culture system. CD34+ NK progenitors were sorted and seeded onto EL08.1D2 stromal cells in the presence of cytokines (24). Using progenitors from 6 donors, we found robust proliferation measured by Ki67 positivity during NK cell development (Fig. 4A). Cells were particularly proliferative at Day 7, when CD34+ progenitors are downregulating CD34 and gaining expression of CD117 (Fig. 4A). We also found increased proliferation at day 21 of culture relative to days 14 and 28, at which time CD117+CD94– NK precursors are becoming CD94+ immature NK cells (Fig. 4A).

**Fig. 4.**
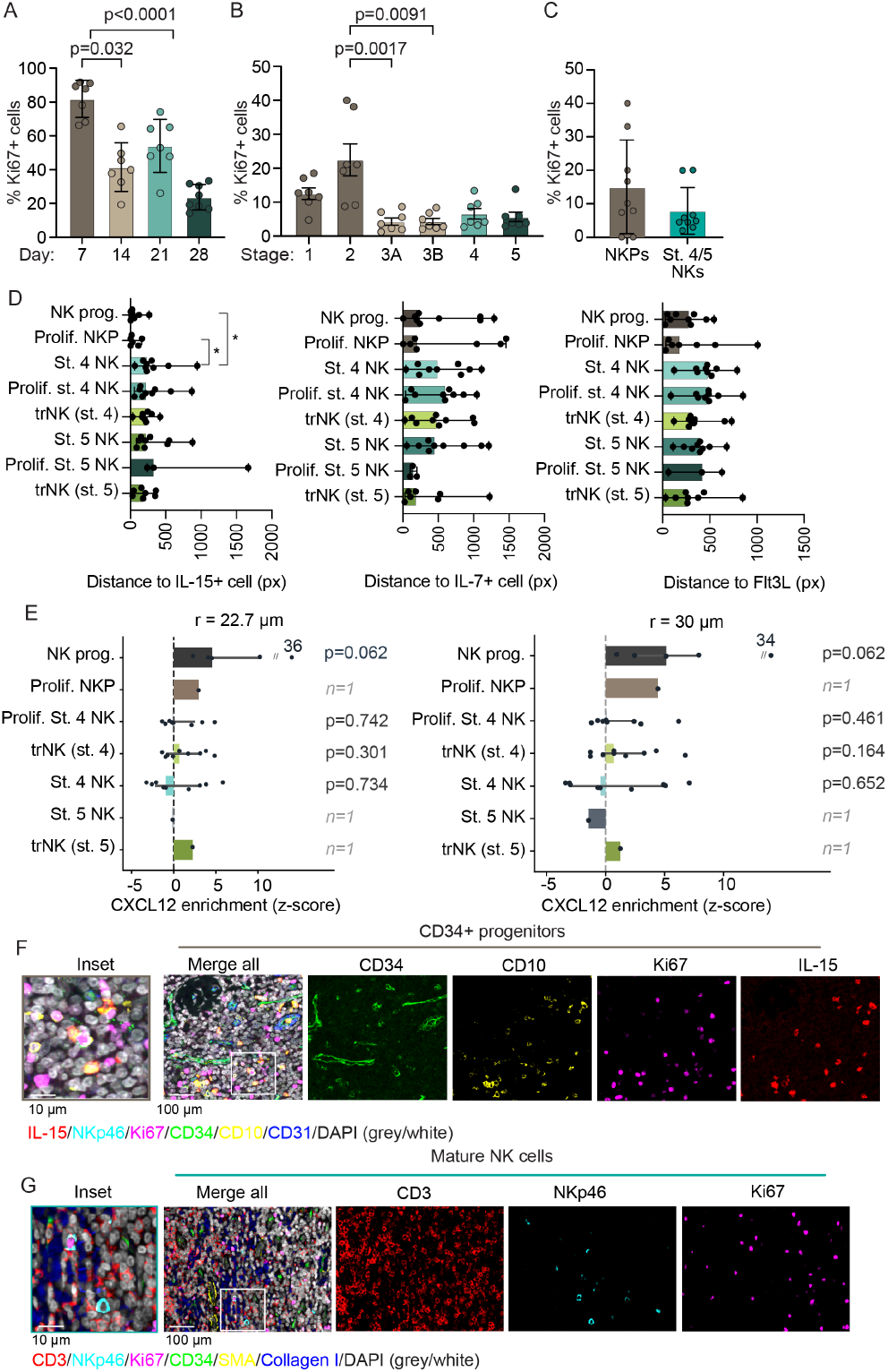
NK cell progenitors proliferate and localize to IL-15+ and CXCL12+ cells in tonsil. A) Frequency of Ki67+ cells during in vitro NK differentiation from CD34+ progenitors co-cultured with EL08.1D2 stromal cells (n = 6 donors). Bars show mean ± SD; dots represent individual donors at each time point. One-way ANOVA with Tukey’s post hoc test; selected pairwise comparisons shown. B) Frequency of Ki67+ cells in NK developmental stages 1–5 by flow cytometry of pediatric tonsil donors (n = 7 donors). Bars show mean ± SD; dots represent individual donors. Kruskal-Wallis with Dunn’s post hoc; selected comparisons shown. C) Frequency of Ki67+ cells in NKPs (stages 1–2) vs. mature NK cells (stages 4–5) by CyCIF (n = 9 donors). Bars show mean ± SD; dots represent individual donors. D) Per-donor median shortest distance (pixels; 0.454 µm/pixel) from each NK developmental subset to the nearest IL-15+, IL-7+, or Flt3L+ cell, by CyCIF (n = 9 donors). Bars show mean across donors; dots represent individual donors. *p < 0.05, paired Wilcoxon vs. NK progenitor; only significant comparisons shown. E) Spatial enrichment of NK developmental subsets for CXCL12+ cells quantified as a permutation-based z-score (observed vs. expected mean CXCL12+ neighbors within 22.7 or 30 µm radius). Each point is one donor; bars show median with IQR. Donor rank was consistent between both radii measured (Wilcoxon p = 0.062, the minimum available value for n=5). One donor (ENT100, z 36/34) is plotted off-scale (//, value annotated). Subsets detected in fewer than 3 donors are shown with hatched bars and an italic n flag and were not tested. F) Representative CyCIF images of CD34+ progenitors in tonsil. Left, high-magnification inset (10 µm scale bar); middle, merged view (100 µm scale bar) showing the inset region (white box). Right, single-channel images for CD34, CD10, Ki67, and IL-15. Merge color key: IL-15 (red), NKp46 (cyan), Ki67 (magenta), CD34 (green), CD10 (yellow), CD31 (blue), DAPI (grey/white). G) Representative CyCIF images of mature NK cells in tonsil. Panel arrangement and scale bars as in (F). Merge color key: CD3 (red), NKp46 (cyan), Ki67 (magenta), CD34 (green), SMA (yellow), Collagen I (blue), DAPI (grey/white).

To validate our in vitro observations, we measured Ki67 expression of NK cell developmental intermediates. Consistent with our in vitro differentiation results, stage 1 and 2 NK cells from tonsil had significantly higher expression of Ki67 than stages 3-5 when measured by flow cytometry (Fig. 4B). These results were further validated using CycIF, where we observed a higher fraction of Ki67+ NK progenitors (stage 1-2) compared to mature NK cells (Fig. 4C). Together, these data confirm previous findings describing proliferation of NK cells after lineage commitment (25) (Fig. 4C).

Having confirmed differences in proliferation between NK subsets, together with NK cell subset localization (Fig. 1) and cell-cell interactions (Fig. 3), we wanted to understand whether NK progenitors are more likely to be interacting with cells expressing cytokines that support NK cell development. Quantification of the median shortest distance was used to characterize interactions between NK cell subsets and cytokine-producing cells including IL-15+, IL-7+, and Flt3L+ cells (Fig. 4D). This quantification revealed that NK progenitors and proliferating NK progenitors were preferentially localized within 30 µm of IL-15+ cells, significantly closer than mature Stage 4 or Stage 5 NK cell subsets, which were typically positioned >200 µm from the nearest IL-15+ cell (Fig. 4D). A similar, but not significant, pattern was observed for Flt3L, where NK progenitors were closer to Flt3L+ cells than mature NK subsets, while no consistent differences across subsets were observed for IL-7.

In mouse bone marrow, developing NK cells are found in proximity to CXCL12 abundant reticular cells that express IL-15 (25). To determine whether a similar niche supports NK cell development in human tonsil, we calculated a permutation-based enrichment z-score for probability of interactions between NK cells and cells expressing CXCL12. We performed this analysis at two radii (22.7 µm and 30 µm). Subsets were included for a given donor only if n >20 cells of that subset were present, and only subsets meeting this threshold in >5 donors were formally tested (see Methods). Consistently, we found that NK progenitors were significantly enriched in proximity to CXCL12+ cells (5/5 donors) (Fig. 4E). Visualization of representative CyCIF images showed NKPs in proximity to IL-15+ cells (Fig. 4F) and NKp46+ NK cells in T cell-rich areas (Fig. 4G).

Together with the proliferation data in panels A–C and the stromal association of NK progenitors observed in Fig. 3, the close spatial relationship of NK progenitors with IL-15+ cells suggests that IL-15 supports a progenitor proliferative niche in tonsil, consistent with its established role in NK cell development. NK progenitors were also enriched in the neighborhood of CXCL12+ cells, positioning the earliest progenitors in tonsil within an environment containing supportive cytokines and chemokine cues. Cell proliferation at the NKP and immature NK cell stages thus appears to be a conserved feature of NK cell development in vitro and in vivo, with IL15 and Flt3L as the proximate cytokine signals and CXCL12 as a candidate spatial signal in tissue.

### Identification and functional validation of a tonsil stromal niche supporting NK cell development

Having localized NK cell developmental subsets to specific tissue domains and identified cytokine signals associated with proliferation, we asked which cell populations constitute this supportive niche. We reasoned that the niche should comprise stromal cells expressing the developmental cytokines (IL-15, IL-7, Flt3L) and adhesion or signaling molecules (DLL1, DLL4, JAG1, JAG2) required for NK cell maturation, and should be spatially positioned to interact with NKPs and developing NK cells in situ.

Nanostring GeoMx analysis of 32 manually defined ROIs from 6 pediatric tonsil sections revealed that parafollicular and interfollicular domains had higher expression of developmental cytokines and ligands than follicles (Fig. 5A). Specifically, we found that DLL4, FLT3L, SCF, IL7, JAG1 and JAG2 transcripts were more highly expressed by cells within interfollicular and/or parafollicular domains (Fig. 5A). IL15 expression was low but detected in the interfollicular and parafollicular domains (Fig. 5A). We visually confirmed these ligands by immunofluorescence, which demonstrated the expression of IL-7 in the parafollicular and subepithelial domains and IL-15 and Flt3L in interfollicular domains (Fig. 5B). As expected, these ligands were found on stromal and endothelial cells, and gp38+ FRCs expressed IL-7 and IL15 in the interfollicular, parafollicular, and subepithelial domains (Fig. 5B). Quantification of the relative fluorescence intensity (Z-scored per cytokine) of CXCL12, Flt3L, IL-15, and IL-7 across cell populations identified by CyCIF showed that stromal, endothelial, and epithelial cells express NK developmentally supportive cytokines, with CXCL12 most highly expressed by endothelial cells and IL-15 by epithelial cells (Fig. 5C).

**Fig. 5.**
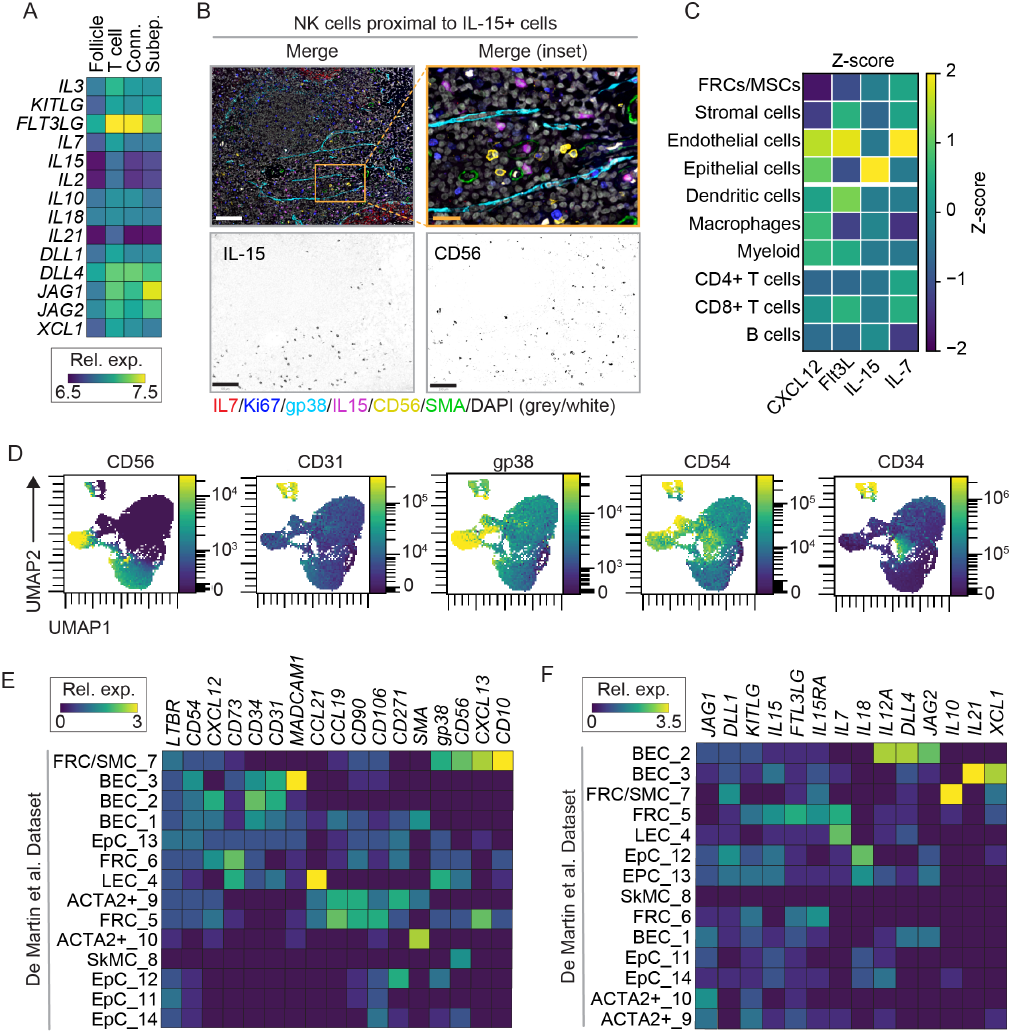
Tonsils harbor cytokine-producing stromal cells that can support NK cell maturation. A) Nanostring GeoMx analysis of select Notch ligands, cytokines, and chemokines. Gene expression data from 32 ROIs from 6 donors was normalized and log2 scaled. B) Proliferating NK cells in proximity to IL-15+ and IL7+ cells in the interfollicular domain. Right image is an enlarged ROI from the image on the left. Greyscale images are shown below to highlight IL-15 and CD56 expression. Representative of 9 ROIs from 9 donors. Scale bar=100 µm (left), 20 µm (right). C) Z-scored mean fluorescence intensity of CXCL12, Flt3L, IL-15, and IL-7 across non-NK cell populations identified by CyCIF (n=9 donors). Z-scores are computed per cytokine (column-wise). D-F) Distance to Flt3L+, IL7+, and IL15+ cells calculated for CD34+ HSPC or CD56+Lin– NK cells. Vertical lines mark percentile as indicated. F) Difference in distance to nearest IL-15+ cell between NKP and mature NK cells. Dashed line indicates mode, n=9 donors. G) UMAP of tonsil (n=3 donors) and bone marrow stromal cells (n=1 donor) showing expression intensity of endothelial and MSC markers. UMAP was calculated using expression data of all markers except for lineage (CD45/CD3/CD14/CD19) and live cell markers. H) Expression of chemokine and cytokines in stromal cell subsets from the dataset published in De Martin et al (61). I) Expression of ligands that function in NK cell development in stromal cells from the dataset published in De Martin et al. (32)

We then used flow cytometry to measure expression of adhesion ligands (ICAM-1, VCAM-1) and markers of mesenchymal (CD90, CD73) or endothelial states (CD31, CD34) (Table S4) (13, 14, 26–29). We included NCAM (CD56), given its role in facilitating human NK cell differentiation (30). Tonsil stromal cells clustered distinctly when visualized by UMAP and we identified a population of tonsil FRCs highly expressing gp38 and CD73 (Fig. 5D). Markers including CD73 and CD90 were expressed on BM stroma and a subset of tonsil stroma, whereas ICAM-1 (CD54) and VCAM-1 (CD106) were more highly expressed on a subset of tonsil stroma than BM stroma. As previously described, a clone of CD56/NCAM (39D5) that specifically recognizes BM stromal cells was detected on a subset of tonsil stroma (31), as was conventionally detected CD56/NCAM (MY31) (Fig. 5D).

Analysis of scRNA-Seq data generated from tonsil stromal cell populations (32) identified two FRC populations (Fig. 5E; FRC_5 and FRC/SMC_7) that reflect the NCAM+ FRCs we found in proximity to NKPs (Fig. 5B) and identified by flow cytometry (Fig. 5D). At the transcript level, IL15, IL7, and FLT3L were expressed by gp38^low^ CD56^low^ CXCL13+ CCL19+ CD106+ CD90+ FRCs (FRC_5), while gp38^low^ CD56^low^ CD34+ CCL21+ lymphatic endothelial cells (LEC_4) expressed IL7 exclusively (Fig. 5F). FRC_5 and FRC/SMC_7 correspond to the FRC/MSC subset and LEC_4 to the endothelial subset in our CyCIF dataset (Fig. 5E). Co-expression of IL15RA and IL15 by FRC_5 cells suggests that they can trans-present IL-15 to developing NK progenitors, whereas blood endothelial cells (BEC_2 and BEC_3) showed low cytokine expression (Fig. 5F). At the protein level, CyCIF confirmed cytokine expression across stromal, endothelial, and epithelial compartments, with endothelial cells showing the highest relative CXCL12 and IL7 and epithelial cells the highest IL-15 (Fig. 5C). Together, these data indicate that multiple stromal and endothelial populations contribute developmentally supportive cytokines to the tonsil NK niche, and visualization of proliferating NK cells adjacent to these populations is consistent with a local proliferative signal (Fig. 5B).

The in situ identification of a candidate stromal niche prompted us to test whether these stromal cells could functionally support NK cell development. We isolated adherent tonsil stromal cells (TSCs) and assessed their expression of CD31, CD54, CD10, and gp38 by flow cytometry (Fig. 6A). Primary stromal cells were largely CD31 negative nonendothelial mesenchymal cells, including fibroblastic reticular cells (Fig. 6A). RNA-Seq confirmed expression of MSC genes including CD90, CD44, and CXCL12, but not endothelial and epithelial genes (e.g. KRT1, MECA79, and CD34; Fig. 6B). TSCs did not retain significant expression of IL-15, IL-7, KITLG (SCF), or FLT3L, suggesting that in vitro support of NK cell lineage commitment by TSCs is mediated by other factors, possibly GAS6 (33), or by other cells that are generated in concert with NK cells in these cultures.

**Fig. 6.**
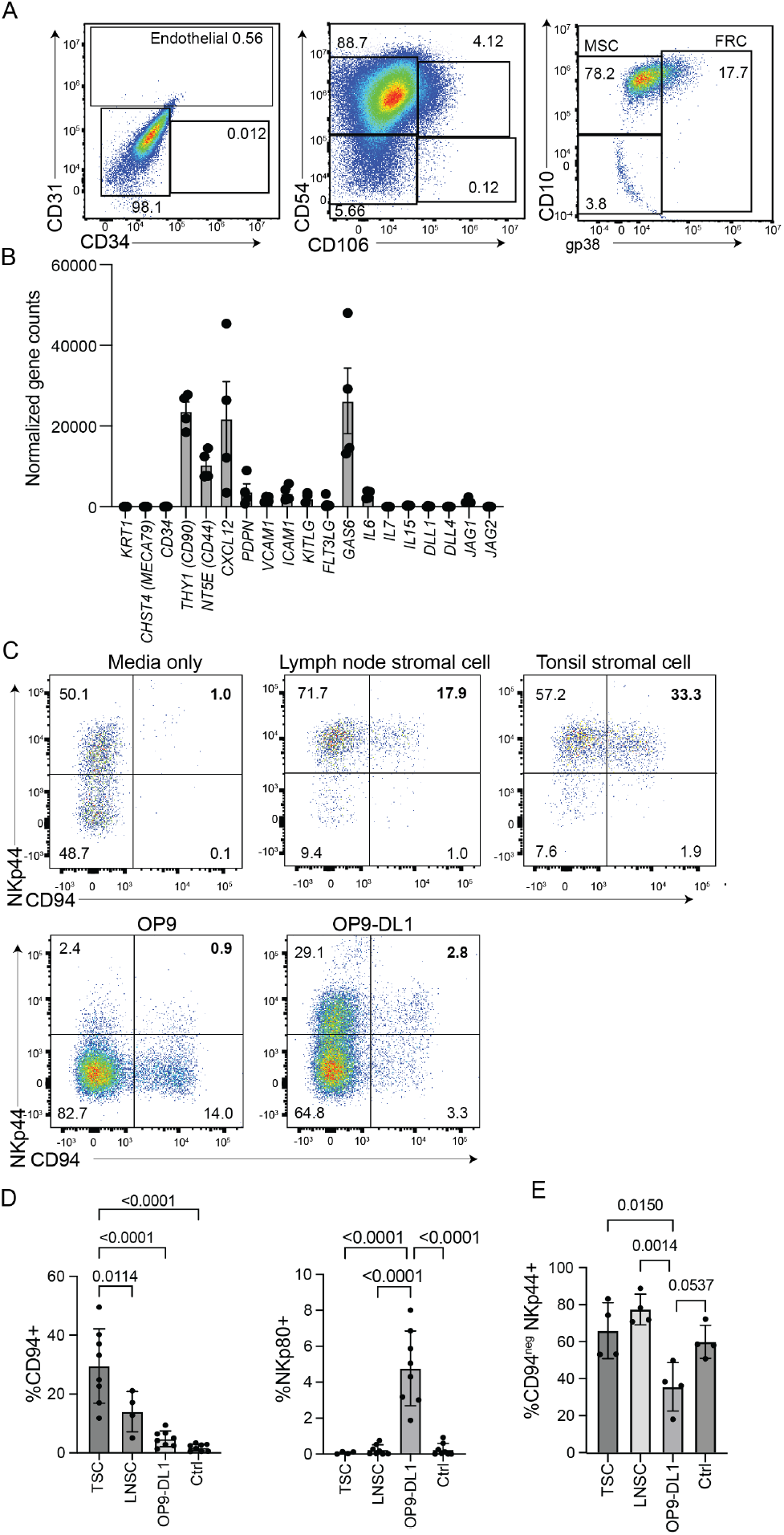
Primary tonsil stromal cells express ligands that support human NK cell development. Stromal cells were isolated from human tonsil or lymph nodes. A) Representative flow cytometric analysis depicting surface receptor expression of in vitro cultured tonsil stromal cells isolated from primary tonsillar tissue acquired from routine pediatric tonsillectomies. B) Bulk RNA-seq was performed on primary tonsil stromal cells from 4 donors. Expression of selected genes encoding ligands known to play a role in NK cell development are shown as normalized gene counts. C) Flow cytometric analysis of NK cell differentiation from CD34+ HSPCs in the presence of cytokines and primary stromal cells or commonly used stromal cell lines. Frequency of NKp44+ CD94+ conventional NK cells (shown in bold font) was measured after 4 weeks of culture. D) Quantification of CD94+ or NKp80+ conventional NK cells (CD94+Lingated on live lymphocytes) after 3 weeks of in vitro differentiation. TSC, tonsil stromal cells; LNSC, lymph node stromal cells; OP9-DL1 or control (media only). n=8 technical and biological replicates. Mean±SD, p-values calculated by one-way ANOVA with multiple comparisons. E) Quantification of non-NK cell ILC subsets (CD94-NKp44+, Lingated on live lymphocytes). TSC, tonsil stromal cells; LNSC, lymph node stromal cells; OP9-DL1 or control (media only). n=4 technical and biological replicates. Mean±SD, p-values calculated by one-way ANOVA with multiple comparisons.

We asked whether tonsil stromal cells could support commitment of Lin–CD34–CD117+ NK progenitors to the conventional NK cell lineage. We compared in vitro NK cell development using tonsil stromal cells (TSCs), lymph node stromal cells (LNSCs), OP9-DL1, OP9, and feeder-free conditions. After 3 weeks of culture in the presence of exogenous IL-7, in vitro derived NK cell populations from each condition were assessed by flow cytometry. Culturing NK progenitors with TSCs led to increased production of CD94+ NK cells after 3 weeks of culture (Fig. 6C, D). As previously reported (16), the presence of Notch ligand on OP9DL1 promoted the upregulation of NKp80 (Fig. 6D). In contrast, LNSCs or TSCs did not produce a significant population of NKp80+ NK cells, suggesting that TSCs lack an essential Notch ligand for later steps of NK cell maturation but can support early NK cell lineage commitment. Similarly, we observed a higher frequency of NKp44+CD94– ILCs in our cultures from TSCs, LNSCs and feeder-free conditions relative to OP9-DL1 (Fig. 6C, E). This demonstrates that tonsil stromal cells can promote NK/ILC cell development from CD34–CD117+ NK progenitors but are not likely the source of Notch ligands required for terminal maturation. Together, these data identify a tonsil stromal population that is spatially positioned and functionally competent to support NK cell development, defining a cellular niche for human NK cell differentiation in secondary lymphoid tissue.

### Local inflammation leads to changes in NK cell frequencies, interactions and localization

We then defined the effects of inflammation on NK cell subset frequency, cellcell interactions, and localization. Inflammation in tonsils used for our study was determined by post-operative pathology reports, which determined the presence of chronic or acute local inflammation at the time of resection (Table S5).

Using CyCIF to assess changes in NK cell subset frequency, we found that non-tissue resident (CD49a and CD103 negative) proliferating and non-proliferating stage 4 NK cells make up a higher percentage of the total cell population in tonsil in the presence of local inflammation (Fig. 7A). We then quantified the frequency of NK cell subsets by flow cytometry and found an increased frequency of NK cells present in inflamed tonsillar tissue relative to noninflamed donors (median 9.8% vs. 4.8%, Fig. 7B). Relative to noninflamed donors, we noted that donors with inflammation present at resection had a significant increase in the frequency of immature and stage 4 NK cells, supporting our CyCIF data (Fig. 7C). To further determine whether inflammation contributed to differential proliferation within each NK cell subset, we compared the frequency of Ki67 positivity within NK cell populations using CyCIF and found that the percentage of Ki67+ cells was increased in NKPs from 2 of 3 donors in response to inflammation (Fig. 7D).

**Fig. 7.**
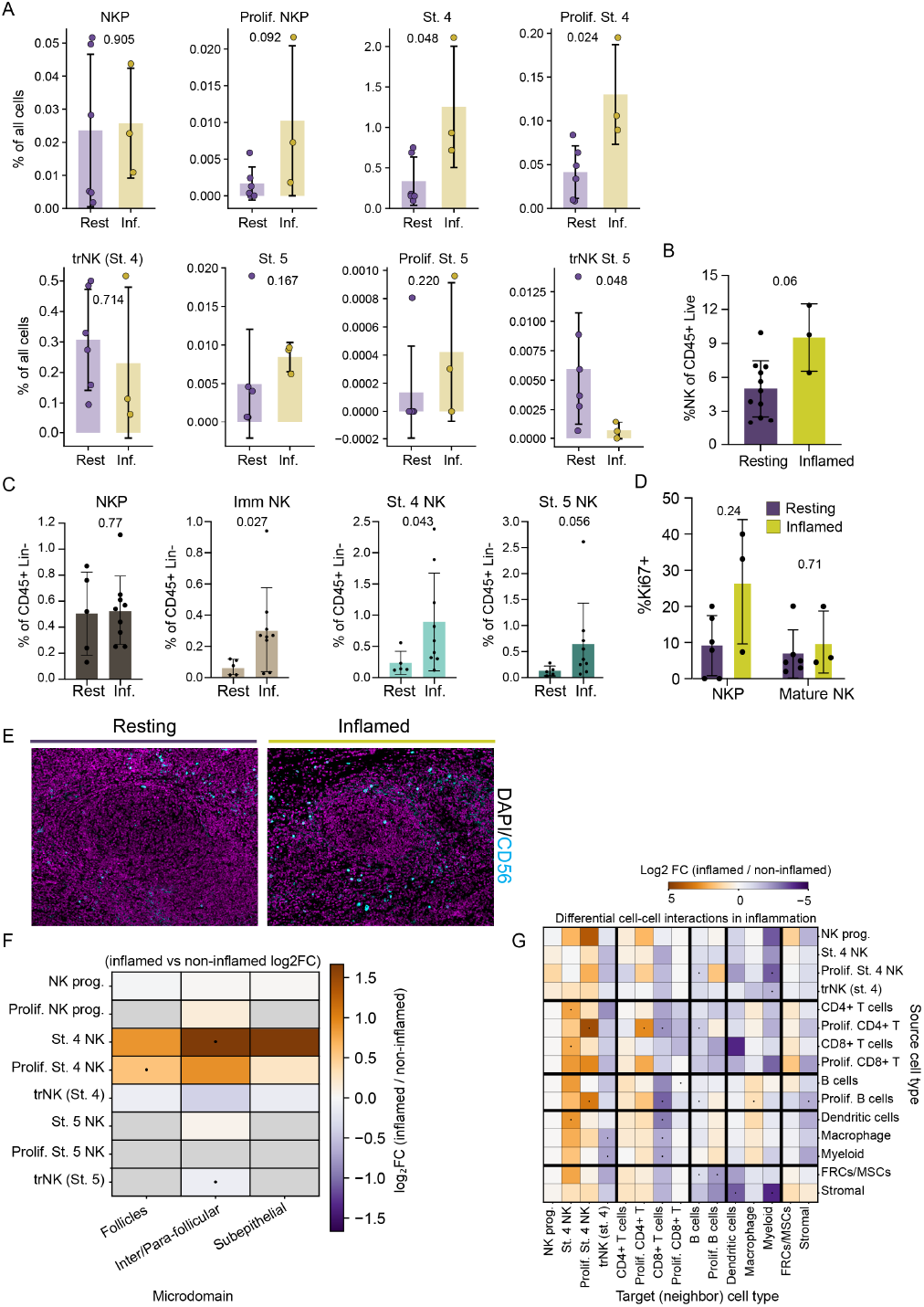
Inflammation affects NK cell subset frequency, localization, and interactions in situ. Frequency of NK cell subsets in inflamed and non-inflamed (resting) tonsil tissues calculated from CyCIF images (n = 6 (resting) or 3 (inflamed)). B) Frequency of NK cells measured by flow cytometry (n = 10 (resting) or 3 (inflamed)). C) Frequency of NK cell subsets measured by flow cytometry (n = 5 (resting) or 8 (inflamed)). D) Frequency of Ki67+ NK cells from CyCIF data (n = 6 (resting) or 3 (inflamed)). E) Representative CycIF image of a follicle and surrounding T cell zone showing CD56 (cyan) and DAPI (magenta) from inflamed and non-inflamed donors. F) Fold change in the frequency of NK cell subsets found in each microdomain identified by CycIF of 3 inflamed and 6 noninflamed tonsil donors. G) Log2 fold change in frequency of NK cell subsets within each microdomain during local inflammation. Frequency calculated as (NK subset cells / all cells in that microdomain) per donor, then compared between inflamed (n=3) and non-inflamed (n=6) groups. Bullet () indicates p<0.05; no pair significant after Benjamini-Hochberg FDR correction. Grey cells indicate below detection threshold (< 20 total cells or < 2 donors with detectable cells). All statistical comparisons were calculated by non-parametric MannWhitney test.

We observed an increase in the abundance of NK cells in and around follicles based on the spatial distribution of CD56+ cells (Fig. 7E). To quantify the change in NK cell localization due to inflammation, we measured changes in frequencies of NK cells found in tissue microdomains (Fig. 7F). We found that proliferating stage 4 NK cells were more abundant within follicles, while non-proliferating stage 4 NK cells were enriched in the inter/parafollicular zone with local inflammation (Fig. 7E, F). We calculated differential interactions between NK cell subsets and other cell types in the presence or absence of inflammation. NK cells did not engage in novel cell-cell interactions in settings of inflammation. Instead, we noted quantitative differences in the likelihood of cell-cell interactions, though none survived correction for multiple comparisons (Fig. 7G). Thus, we found that NK cells make up a larger proportion of CD45+CD3–CD14–CD19– cells in inflamed tonsil, with non-tissue resident stage 4 NK cells contributing the most consistent increases in abundance through their proliferation and localization.

## Discussion

NK cell maturation in tonsil is thought to begin with a circulating CD34+ progenitor analogous to a bone marrow-derived tissue seeding lymphocyte precursor that enters tissue from peripheral blood (1, 7, 8, 33, 34). Although the signals that mediate homing of circulating CD34+ NKPs to SLT have not been defined, expression of CXCR4 and CCR7, which can interact with CXCL12 and CCL21 respectively, suggests that progenitors enter via HEV or blood vessels expressing these chemokine ligands (25, 35, 36). This observation is additionally supported by our data showing proximity of precursors to blood vessels within the interfollicular domain. How, or if, precursors are targeted to thymus or SLT, and similarly whether lineage fate is encoded before their recruitment or occurs solely from stochastic signals in the environment, remains to be determined for human NK and T cell precursors (37).

We identified CD34+CD45+CD122+ progenitors expressing CD49d, the integrin alpha chain associated with integrin β7, in the interfollicular and parafollicular domain adjacent to stromal cells, including FRCs. While we did not explicitly test CD45RA expression, the expression of CD45 and CD49d suggests that these cells are analogous to previously described CD34+CD45RA+integrin β7^bright^ precursors (8). Direct interaction with stromal cells improves the efficiency of NK cell maturation from CD34+ progenitors in vitro, suggesting that interactions between NK cell subsets and stromal cells or FRCs similarly supports early stages of NK cell development in vivo (24, 38–40). The use of tonsil-derived stromal cells for in vitro NK cell differentiation further demonstrates that secondary lymphoid tissue contains cells that promote the production of mature NK cells from circulating NK progenitors. Consistent with this, stage 1–2 NKPs, but not mature subsets, were enriched within CXCL12-rich neighborhoods of the interfollicular/parafollicular stroma (Fig. 4E), where endothelial cells were the predominant CXCL12 source (Fig. 5C), and lay in closer proximity to IL-15+ cells than mature subsets. This perivascular CXCL12+ niche positions progenitors to receive maturation signals locally and links their entry via CXCL12+ HEVs to subsequent stromal residence and proliferation. While CD34+ NKPs are rare in tonsil, our flow cytometry and imaging data show they are the most proliferative NK cell developmental subset, reflective of their proximity to cytokine-producing stromal and endothelial cells. The expression of high affinity IL-2 receptor on NKPs in tonsil suggests that they also can respond to IL2 from T cells, like mature NK cell populations (8, 11). As such, we propose that the trafficking and subsequent proliferation of rare tissue-seeding progenitors can provide a larger pool of stage 3 NK cell precursors that can continue through maturation to become mature ILC or NK subsets following additional differentiation and maturation signals (4).

The localization of NKPs to the parafollicular and interfollicular domains, where FRCs are found, may also be directed in part by CXCL12 and CCL21, as stage 1-2 NKPs expressed CXCR4 and CCR7 and were in proximity to CXCL12+ and CCL21+ cells. While CCR7 was poorly detected by flow cytometry on most NK cell subsets, its gene expression on stage 3 NK cells suggests that we detected this NKP by CyCIF in tissue, and previous studies have reported responsiveness of CD56+CD16– cells to CCR7 ligands (36, 41). CXCR5 expression on NKPs could also direct them towards the follicle, but more likely is important for trafficking into tonsil via CXCL13+ HEVs (28). As NK cells progress though maturation, they move away from the interfollicular domain to the CCL20high parafollicular T cell rich zone and downregulate CCR6. This observation confirms previous observations of NK cells in the parafollicular microdomain where they can respond to IL-2 from T cells (11). Most proliferating stage 4 and 5 NK cells are not in proximity to IL-7, IL-15, or Flt3L expressing cells, suggesting that other factors, including IL2, can promote mature NK cell proliferation in situ (11). In addition, the increased frequency of interactions between mature NK cells and CD4+ T cells underscores the role of NK cells in assisting in the coordination of adaptive immunity (42–44). Finally, while these data identify certain correlations between chemokine receptors and ligands, it should be noted that trafficking in tissue is complex and directed by chemokine signaling, cell adhesion, and physical properties of tissue.

Within the interfollicular/parafollicular domains, trNK and non-trNK cell subsets occupy unique spaces based on their distinct cell-cell interactions and proximity to CXCL12, CCL19, and CCL21. trNK cells have more interactions with macrophage and other myeloid lineage cells than non-tissue resident NK cells, suggesting trNK populations are more likely to be generating or receiving signals from myeloid cells. Non-tissue resident NK cells, although they can interact with myeloid cells, are observed to preferentially interact with T cells, suggesting they are modulating T cell responses and elimination (45). Moreover, we show that these interactions become more prevalent in the presence of local inflammation, further indicating their importance in coordinating innate and adaptive immune responses.

In summary, this resource is a comprehensive analysis of human NK cell developmental subsets in situ that provides insights into the function, developmental niche, and trafficking of NK cell subsets. Our observations support a model in which circulating progenitor cells enter the tonsil via HEVs lined with CXCL12+ and CCL21+ endothelial cells in the interfollicular domain. There, NKPs reside within a cytokinerich and CXCL12+ stromal niche that induces proliferation and further maturation. As progenitors mature into trNK and non-tissue resident stage 4 and 5 NK cells, they relocate to the parafollicular and subepithelial domains and engage T cell subsets. Local inflammation causes NK cells to become more proliferative and abundant in the parafollicular domain, where non-tissue resident NK cells interact with T cells and trNK cells interact with both T cells and myeloid cells. Together, our study defines a road map for human NK cell development at steady state and under inflammatory conditions.

## Methods

### Tissue acquisition and processing

Matched peripheral blood and tonsil tissue were collected from pediatric patients undergoing routine tonsillectomies at Columbia University Irving Medical Center. All tissue and blood samples were collected in accordance with the Declaration of Helsinki, with written and informed consent from all participants under the guidance of the Institutional Review Board of Columbia University. Post-operative pathology reports were used to determine whether local inflammation was present. Tonsil tissue samples were split in two sections to make single cell suspensions and formaldehyde fixed paraffin embedded (FFPE) tissue slides. To produce single cell suspensions from tonsil, specimens were placed into a petri dish with sterile PBS and manually dissociated by mincing the tissue into a cell suspension which was then filtered through a 40 µm cell strainer. Samples were then centrifuged at 1200 rpm for 7 minutes and resuspended in PBS followed by another round of centrifugation. Tonsil cell suspensions were resuspended in heat inactivated FBS with 10% DMSO to cryopreserve for flow cytometric analysis. The remaining tonsil specimens were fixed in 10% paraformaldehyde for 24-48 hours then placed in 70% ethanol. Fixed specimens were given to the Herbert Irving Comprehensive Cancer Center Histology Service and Tumor Banking core for paraffin embedding, tissue sectioning, and slide preparation. 3 µm sections were prepared for imaging mass cytometry and high-resolution microscopy, and 10 µm sections for spatial transcriptomics.

Blood from pediatric tonsil donors was collected by venipuncture on the day of their tonsillectomy. Whole blood was diluted with equal parts of PBS and layered on FicollPaque density gradient followed by centrifugation at 2,400 rpm for 20 mins with no brake and slow acceleration. The lymphocyte layer was collected from gradient interface and washed with equal parts of sterile PBS then centrifuged at 1,200 rpm for 5 mins. Collected cells were resuspended in HI FBS with 10% DMSO at 1-5 x 10^6^ cells/mL and cryopreserved for flow cytometric analysis.

### Spatial transcriptomics

FFPE tissue was prepared into 10 µm sections and sent to Nanostring for GeoMx RNA assay data acquisition on a GeoMx Digital Spatial Analyzer (DSP). 35 domains of interest were chosen based on tonsillar microarchitecture using tissue sections from 6 pediatric donors (4 noninflamed and 2 inflamed). Spatial transcriptomic data was normalized using Q3 normalization after BioQC and filtering using GeoMx DSP Analysis Suite (version 2.5.1.145). Heatmaps were produced using ClustVis (46) and Heatmapper (47).

### Flow cytometry and analysis

Cryopreserved tonsil cell suspensions and PBMCs were thawed and resuspended in sterile PBS with 10% FBS. Before immunostaining, antibodies specific to extracellular proteins were diluted in PBS with 10% FBS (Table S2-4). Cells were stained for extracellular markers for 30 mins at room temperature (RT) then washed. If required, cells were then fixed and permeabilized using eBioscience FoxP3 buffer (catalog no. 00-5523-00; ThermoFisher) then immunostained for intracellular markers for 30 mins at RT.

A Novocyte Penteon Cell Analyzer was used to acquire data, which was then exported to FlowJo (BD Biosciences) or OMIQ (Dotmatics) for downstream analysis. For defining chemokine receptor expression of NK cell developmental subsets, we identified NK cell developmental subsets using single-cell analysis in OMIQ. First, data were arcsinh transformed and live single CD45+lin– cells were gated. FlowAI (47) was applied to remove low quality cells and FlowSOM (48, 49) based on all the markers of the backbone panel was applied to remove any residual lineage positive dead cells and innate lymphoid cells (CD127+NKp44+, CD127+CD103+, and CD127+CD294+). Chemokine receptor positive gating on NK cell subsets was determined by fluorescence minus one control. For flow cytometric analysis of human stromal cells from bone marrow and tonsil, data were arcsinh transformed then cells were gated to remove lineage positive (CD14+, CD19+, CD45+, and/or CD3+) and dead cells before performing UMAP dimensional reduction analysis using all markers.

### CyCIF microscopy

FFPE tonsil sections were stored at 4C and brought to room temperature prior to dewaxing with three 10-minute Xylene (catalog no. X5–1; Fisher Scientific) washes. Next, tissue was rehydrated using serial washes in 100% and 95% ethanol (catalog no. BP2818100; Fisher Scientific) for 10-minutes, then in 70% and 35% ethanol for 5 minutes. Excess ethanol was washed out with water. Heat-activated antigen retrieval was carried out on rehydrated tissue sections by incubating slides in pre-warmed Agilent Dako pH6 antigen retrieval solution (catalog no. S2367; Agilent) for 20-minutes in a vegetable steamer (Black Decker), cooled to room temperature, and washed with PBS. For Cy-CIF, tissue sections were incubated with 1 mg/mL borohydride in deionized water for 10 minutes to decrease background fluorescence then washed three times with PBS. Samples were permeabilized with 0.2% Triton X-100 in PBS for 30 minutes and washed with PBS three times. After permeabilization, tissue samples were incubated with ThermoFisher Super Blocking Buffer (catalog no. 37580; Thermo Fisher) for 1 hour. Antibodies for immunostaining were diluted in antibody diluent buffer [PBS/10% Bovine Serum Albumin/0.01% Triton X-100]. Immunostaining of tissue sections with unconjugated antibodies for high-resolution immunofluorescence microscopy was done overnight at 4C in a humidity chamber.

For immunofluorescence microscopy, unbound antibodies were washed with antibody diluent buffer four times after immunostaining. Tissue sections were incubated with secondary antibodies at a dilution of 1:200 for 1 hour at room temperature followed by four washes with antibody diluent buffer and two washes with PBS. Washing buffer was removed from tissue prior to adding ProLong Glass Antifade Mountant with NucBlue Stain (catalog no. P36985; ThermoFisher) and a 1.5 coverslip (catalog no. 72204-03; Electron Microscopy Services). Samples were cured overnight at room temperature prior to imaging with a Zeiss CellDiscoverer-7 using a 20x/0.7 NA objective and an optical magnification of 0.5x. For CyCIF microscopy, tissue was treated with Vector TrueVIEW Autofluorescence Quenching Kit (catalog no. SP-8400; Vector Labs) prior to mounting. After washing excess TrueView reagents, tissue sections were mounted using VectaShield Antifade Mounting Medium with DAPI (catalog no. H-1200; Vector Laboratories, Inc) and a 1.5 coverslip (catalog no. 72204-03; Electron Microscopy Services). Regions of interest were imaged with a 20% overlap using a Zeiss CellDiscoverer 7 using a 20X/0.7 or 20X/0.95 NA objective and an optical magnification of 0.5X and coordinates were saved for subsequent imaging. This resulted in images with a micron per pixel unit of 0.454 microns/pixel. Upon completion of imaging, coverslips were gently removed by placing slides in a Coplin Jar with PBS. To strip antibodies, tissue slides were incubated in freshly prepared stripping buffer (20 mL 10% SDS/0.8 mL beta-mercaptoethanol/12.5 mL 0.5 M Tris-HCl (pH 6.8)/67.5 mL DI H2O) for 30 mins at 65C then washed with water for 15 mins. Slides were then blocked and re-stained with primary and secondary antibodies prior to re-imaging as described above. 43 unique markers (68 total) were captured by CyCIF (Table S1) over the course of 23 cycles of staining and imaging.

### CyCIF data analysis

For CyCIF image preprocessing, Illumination correction was applied using BaSiC darkfield and flat-field profiles (23). Stitching of tiles and alignment of channels was done using Ashlar with a barrel correction [value used: 7.943e-09]. Background subtraction (Backsub) module was applied to reduce autofluorescence in all channels except for collagen, CD10, CD3, and IL-15. These channels were omitted from background subtraction as the signal to noise ratio was not high enough. Additionally, 21 markers were omitted from downstream analysis based on poor staining represented by low signal to noise ratio and/or nonspecific staining (Table S1). Nuclei-based cell segmentation was performed based on DAPI nuclear staining from first cycle using HALO AI (Indica Labs) or Mesmer (21, 23). CD20, CD127, CD45, CD10, and CD3 were used to define cell membrane boundaries for Mesmer membrane-based cell segmentation. Single-cell data was extracted using McQuant. All packages, except Ashlar, were applied using the McMicro pipeline (23). Subsequent single-cell analysis was performed with SCIMAP (50) (https://scimap.xyz/). For cell phenotyping, CyCIF fluorescence intensity data was first scaled based on manual gating which defined positive and negative intensity thresholds for each marker. The scaled data was then used to define populations based on a phenotype workflow which provided the marker expression associated with each population. NK cells were identified using the strategy in Figure S1A after excluding lineage positive cells (Lineage = CD3, CD20 CD31, smooth muscle actin (SMA), gp38, pankeratin, collagen type 1). For quantifying cell abundance using SCIMAP, tonsil domains were annotated by two complementary approaches. First, domains were manually annotated based on the marker intensity of CD3, CD20, CD10, collagen type 1, smooth muscle actin, pan-keratin, and gp38. Second, a HALO AI (Indica Labs) classifier was trained by manually defining representative regions of each domain; the trained network then considered all marker intensities to assign tissue domains across each section and to quantify CXCL12, CXCL13, CCL19, CCL20, and CCL21 expression (Fig. 1A, 1H). Domain assignments from the trained classifier were concordant with manual annotation. SCIMAP was also used to measure log2-fold differences in cell-type abundance, perform spatial co-occurrence analysis, and calculate distances between cell types.

Spatial co-occurrence analysis shown in Figure 3B was calculated using the radius method to measure all cells within a 50-pixel (22.7 µm) radius from the center of each cell to get the probability and significance of cells colocalizing. The shortest-distance function in SCIMAP was used to compute, for each cell, the distance (µm) to the nearest cell of each target type; these per-cell distances were summarized as the median per donor for analysis and display (Fig. 2D, 2E; Fig. 4D). Microscopy data was visualized using Fiji (51, 52), QuPath (53), and Indica Labs HALO AI software. Prism 9 (GraphPad) and Seaborn [Python version 3.9] was used to plot and statistically analyze CyCIF data (54, 55).

To quantify the spatial association between NK developmental subsets and CXCL12 cells within the inter/para-follicular zone, we computed a permutation-based enrichment z-score for each subset in each donor. For a given subset, we counted the observed mean number of CXCL12 cells within a fixed radius of each subset cell. A null distribution was generated by randomly permuting CXCL12 labels across all cells in the region 200 times per donor while preserving the total number of CXCL12 cells and recomputing the mean neighbor count for each permutation. The enrichment z-score was defined as (observed mean_null) / SD_null, such that z = 0 indicates no enrichment relative to chance and positive values indicate enrichment of CXCL12 cells in the local neighborhood. The analysis was performed at two neighborhood radii, 22.7 µm (50 px) and 30 µm, to confirm robustness to the choice of radius. Subsets were included for a given donor only where >20 cells of that subset were present to ensure stable neighbor estimates; subsets meeting this threshold in fewer than three donors were not formally tested and are indicated individually as an n flag. For subsets present in >5 donors, enrichment across donors was assessed against the null hypothesis of z = 0 using a two-sided Wilcoxon signed-rank test; for n = 5 the minimum attainable two-sided p-value is 0.062. NK progenitors met the inclusion threshold in 5 donors and were enriched for CXCL12 neighbors in all 5 (5/5 donors positive at both radii; donor rank order across radii was identical). Data are shown as the median across donors with interquartile range; individual donors are plotted as points. One donor (ENT100) showed markedly higher enrichment (z = 36 at 22.7 µm; 34 at 30 µm) and is plotted off-scale with an axis break (//); enrichment in NK progenitors remained consistent after excluding this donor.

Inflammation status for each donor was assigned from postoperative pathology reports (Table S5). For each CyCIF donor, the frequency of each NK developmental subset was calculated as a percentage of all segmented cells in the tissue (subset cells / total cells per donor), rather than as a percentage of total NK cells. The independent (per–total cell) denominator was used because subset-of-NK frequencies are compositional: an increase in one subset necessarily lowers the apparent frequency of others, which can produce artifactual decreases (e.g., in tissue-resident Stage 4 NK cells) that are not present when each subset is quantified against the full cell population. Subset frequencies were compared between inflamed (n = 3) and non-inflamed (n = 6) donors by two-sided Mann-Whitney test, with each donor contributing a single value. Subsets were quantified using the residency and proliferation gates defined in Fig. S1A.

To assess inflammation-associated changes in NK cell localization, each cell was assigned to one of three tissue microdomains (follicles, inter/parafollicular, or subepithelial) based on the neighborhood domain annotations described above. Within each microdomain, the frequency of each NK developmental subset was calculated per donor as a fraction of all cells in that microdomain. Subset–microdomain combinations were considered detectable only where 20 cells of the subset were present and 2 donors had detectable cells in that microdomain; combinations below this threshold were not tested and are shown in grey. For detectable combinations, frequencies were compared between inflamed (n = 3) and non-inflamed (n = 6) donors by two-sided Mann-Whitney test, and the inflamed-versus-non-inflamed difference is reported as a log2 fold change of group means.

Pairwise spatial co-occurrence between NK developmental subsets and other tonsil cell types was computed per donor using the 50-pixel (22.7 µm) radius method described above. For each source–target cell-type pair, the per-donor interaction likelihood was compared between inflamed (n = 3) and non-inflamed (n = 6) donors and expressed as a log2 fold change (inflamed / non-inflamed). Statistical significance was assessed by two-sided Mann-Whitney test, and p-values were corrected for multiple comparisons across all tested pairs using the Benjamini-Hochberg false discovery rate procedure. Pairs reaching uncorrected p < 0.05 are indicated; no pair remained significant after FDR correction. Rare populations with insufficient cells for stable per-donor estimates were excluded prior to analysis.

Ki67 staining intensity in the 9-donor CyCIF dataset was provided as log1p-transformed, min-max–normalized values (range 0–ln2). A cell was classified as Ki67-positive if its normalized Ki67 value exceeded 0.42, corresponding to approximately the 92nd percentile of the global intensity distribution; this threshold was selected empirically to reproduce the positivity rates of the original analysis. For each donor, the percentage of Ki67-positive cells was calculated within NK progenitors (NK progenitor and proliferating NKP phenotypes) and within mature NK cells (Stage 4, Stage 5, and their tissue-resident and proliferating subsets). Percentages were compared between inflamed (n = 3) and non-inflamed (n = 6) donors by two-sided Mann-Whitney test. NK progenitor estimates are based on small per-donor cell counts (4–136 cells per donor) and should be interpreted as trends. The percentage of Ki67-positive cells was moderately, but not significantly, increased in NK progenitors in inflammation (p = 0.15) and was unchanged in mature NK cells (p = 0.71).

### In vitro NK cell differentiation

Primary human tonsil stromal cells were maintained at 37C in culture flasks with 90% DMEM (Life Technologies), 20% HI-FBS (Atlanta Biologicals), and 1% penicillin/streptomycin (100 U/mL, Life Technologies). Primary human tonsil stromal cells were isolated from pediatric donors undergoing routine tonsillectomies at Ohio State University Medical Center. Tonsil-derived stromal cells were passaged every 2-3 days and kept in culture for a maximum of 3 weeks.

Leukocyte enriched blood products were acquired from the New York Blood Center,and peripheral blood was collected from healthy donors at CUIMC as a source for CD34+ progenitors. Peripheral blood mononuclear cells were separated from blood samples by Ficoll-Paque density gradient (Cytiva; catalog no. 17144003). To enrich NK cell developmental subsets and CD34+ progenitors from peripheral blood samples, StemCell Rosette Sep negative selection cocktails were used to treat blood products prior to Ficoll-Paque density gradient separation (NK cell enrichment, STEMCELL Technologies; catalog no. 15065; Hematopoietic stem cell enrichment, STEMCELL Technologies; catalog no. 15066).

Enriched blood NK cell developmental subsets were stained anti-CD34 PE (Biolegend, catalog no. 343606), ZombieNIR (Biolegend, catalog no. 423106), and Lineage cocktail (CD3, CD14, and CD19; antibody information in Table S3) for FACS sorting using a BD Aria II cytometer with an 85-µm nozzle or a Sony MA900 Cell Sorter with a 100 µm sorting chip.

In vitro NK cell differentiation for the purpose of measuring proliferation (Fig. 4A) was adapted from Miller et al, 2005 (56). EL08.1D2 and tonsil derived stromal cells were plated into pre-gelatinized, tissue culture treated, flat bottom 96-well plates (Genesee Scientific; catalog no. 25-109) 24-48 hours prior to the start of the differentiation assay. Once stromal cells reached confluency, they were irradiated (30 Gy). CD34+ NK cell progenitors (1-2 x 103) were cultured with stromal cell lines for a total of 3 weeks at 37C in differentiation media containing Ham F12 media plus DMEM (1:2) with 20% heat-inactivated human AB serum, ethanolamine (50 µM), ascorbic acid (20 mg/ml), sodium selenite (5 µg/ml), β-mercaptoethanol (24 µM), and penicillin/streptomycin (100 U/ml) in the presence of IL-15 (5 ng/ml), IL-3 (5 ng/ml), IL-7 (20 ng/ml), stem cell factor (20 ng/ml), and Flt3L (10 ng/ml) (all cytokines from PeproTech). Media exchanges were carried out after each week of culture. Cells were collected after each week of culture and assessed by flow cytometric analysis using the flow panel in Table S3 with the addition of anti-granzyme B BV421 (Biolegend; catalog no. 515408), anti-Ki67 PE (Biolegend; catalog no. 350504), anti-TBET PE-Cy7 (Biolegend; catalog no 644824), and anti-EOMES APC (Invitrogen; catalog no. 50-4877-42). Flow cytometric data was acquired on an Agilent Novocyte Penteon (5-laser) with NovoExpress software and exported for downstream analysis using FlowJo V10 (BD Biosciences).

In vitro NK cell differentiation carried out to test the supportive characteristics of TSC and LNSCs relative to OP9 and OP9-DL1 were carried out as described in Nalin et al., 2020 (16). In brief, FACS sorted CD34+ NKP were initially seeded into culture at 1000 cells per well and with IL-7 (10 ng/ml; Miltenyi) for a total of 14–28 days in 200 µl culture media per well in 96-well flat-bottom plates (TrueLine). FACS sorting, culture media recipe, and phenotypic analysis was carried out as previously described (16).

### Quantitative real time-PCR and flow cytometric analysis of tonsil derived stromal cells

mRNA was isolated and purified using the Total RNA Purification Kit (Norgen Biotek) from FACS-purified primary tonsil-derived stromal cell population according to manufacturer’s instructions and cDNA was synthesized using Superscript IV VILO Master Mix (Thermo Fisher Scientific). Standard quantitative realtime RT-PCR reactions were performed on a Viia7 Real-Time PCR System (Life Technologies) using PowerUp SYBR Green Master Mix (Thermo Fisher Scientific) and primer sequences obtained from a published report (57). Gene expression was normalized to the 18S mRNA internal control: ΔCt = Ct(gene of interest) Ct(18S). Relative mRNA expression for each gene was calculated as 2^(−DCt)^. Gene expression was measured using RT-PCR. For the RT-PCR data, the raw Ct values of the target genes were first normalized to the Ct values of the 18S internal controls.

Flow cytometric analysis of tonsil derived stromal cells was carried out on the adherent fraction from a single cell suspension of tonsil cells. Adherent cells were cultured in alphaMEM (Gibco) with 10% heat inactivated FBS (Gibco) and 1% Pen/Strep (Gibco) at 37C and 5% CO2. Adherent cells were collected and stained with myeloid and lymphocyte lineage markers (CD3, CD14, CD19, and CD45; details for antibodies found in Table S3), plus stromal cell markers including CD31, CD34, CD56, CD106, gp38, and CD10 (Biolegend). Data was collected on a Agilent Novocyte Penteon Cytometer and analyzed using FlowJo.

## ACKNOWLEDGEMENTS

The authors wish to thank research coordinators Evelyn Hernandez and Carlos Aguilar Breton for their assistance with the acquisition of tonsillar tissue from healthy donors. We would like to acknowledge the Columbia Stem Cell Initiative Flow Cytometry Core Facility at Columbia University Irving Medical Center, under the leadership of Michael Kissner, which was used to perform all flow cytometric analysis and cell sorting. We would also like to thank the Herbert Irving Comprehensive Cancer Center Histology Services and Tumor Banking core at Columbia University Irving Medical Center for assistance with tissue embedding and sectioning. We thank Dr. Julie Canman for assistance with figure preparation.

B.I. is a consultant for or received honoraria from Volastra Therapeutics, Johnson & Johnson/Janssen, Novartis, Eisai, AstraZeneca and Merck, and has received research funding to Columbia University from Agenus, Alkermes, Arcus Biosciences, Checkmate Pharmaceuticals, Compugen, Immunocore, and Synthekine. A.G.F. is a consultant for ImmunoVec, Inc and Immunebridge, Inc.

