## Supplemental Material for "Mapping human natural killer cell development in tonsil"

A

| Natural killer (NK) cell subtypes (tonsil) | Identification* |
|---|---|
| NK progenitor (NKP) | LIN- CD45+ CD34+ CD122+ CD127+ CD49d+ |
| Proliferating NK progenitor (NKP) | LIN- CD45+ CD34+ CD122+ CD127+ CD49d+ Ki67+ |
| Immature NK | LIN- CD45+ CD56+ CD127+ GZMK- GZMB- |
| CD56+/NKp46+ NK | LIN- CD45+ CD56+ NKp46+ |
| NK (stage 4) | LIN- CD45+ CD56+ CD127+/- GZMK+/- GZMB- |
| Proliferating NK (stage 4) | LIN- CD45+ CD56+ CD127+/- GZMK+/- GZMB- Ki67+ |
| NK (stage 5) | LIN- CD45+ CD56+ CD127- GZMB+ CD57- |
| Proliferating NK (stage 5) | LIN- CD45+ CD56+ CD127- GZMB+ CD57- Ki67+ |
| trNK (stage 4) | LIN- CD45+ CD56+ CD127+/- GZMK+/- GZMB- CD49a+ CD69+ |
| Proliferating trNK (stage 4) | LIN- CD45+ CD56+ CD127+/- GZMK+/- GZMB- CD49a+ CD69+ Ki67+ |
| trNK (stage 5) | LIN- CD45+ CD56+ CD127- GZMB+ CD57- CD49a+ CD69+ |
| Proliferating trNK (stage 5) | LIN- CD45+ CD56+ CD127- GZMB+ CD57- CD49a+ CD69+ Ki67+ |

NOTE: All cells CD45+. *Lineage negative (LIN-) = CD3- CD13- CD14- CD15- CD19- CD10-

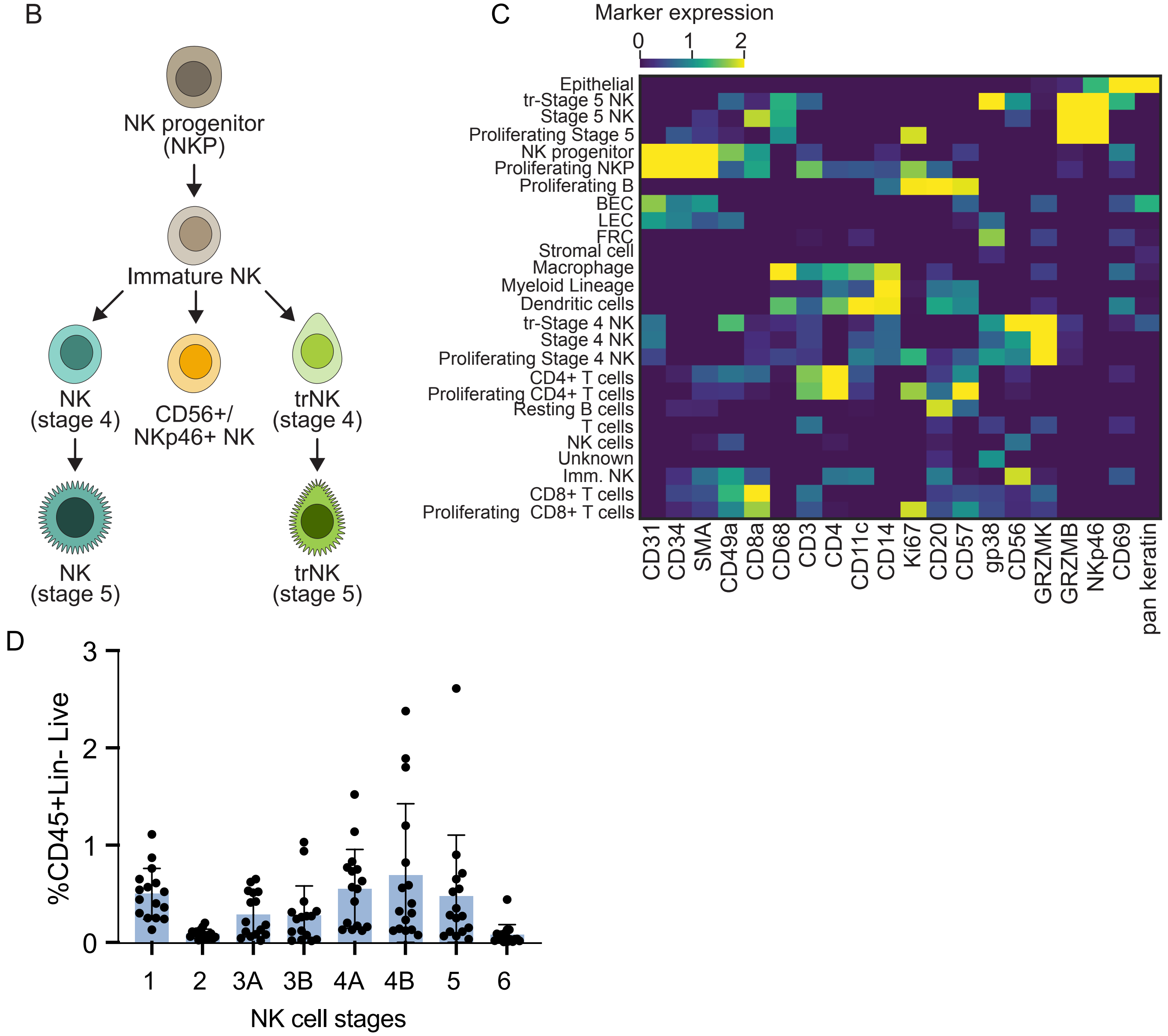

Figure S1: A) Gating strategy used to identify NK cell populations by CyCIF. B) Schematic depicting NK developmental trajectory found in tonsil. C) Relative expression data of phenotypic markers used to identify tonsil cell populations from CyCIF (n=9 donors; 9 ROIs). D) Quantification of the proportion of NK cell subsets within tonsil CD45+CD3–CD19–CD14–Live cells as measured by flow cytometry (n=15 donors).

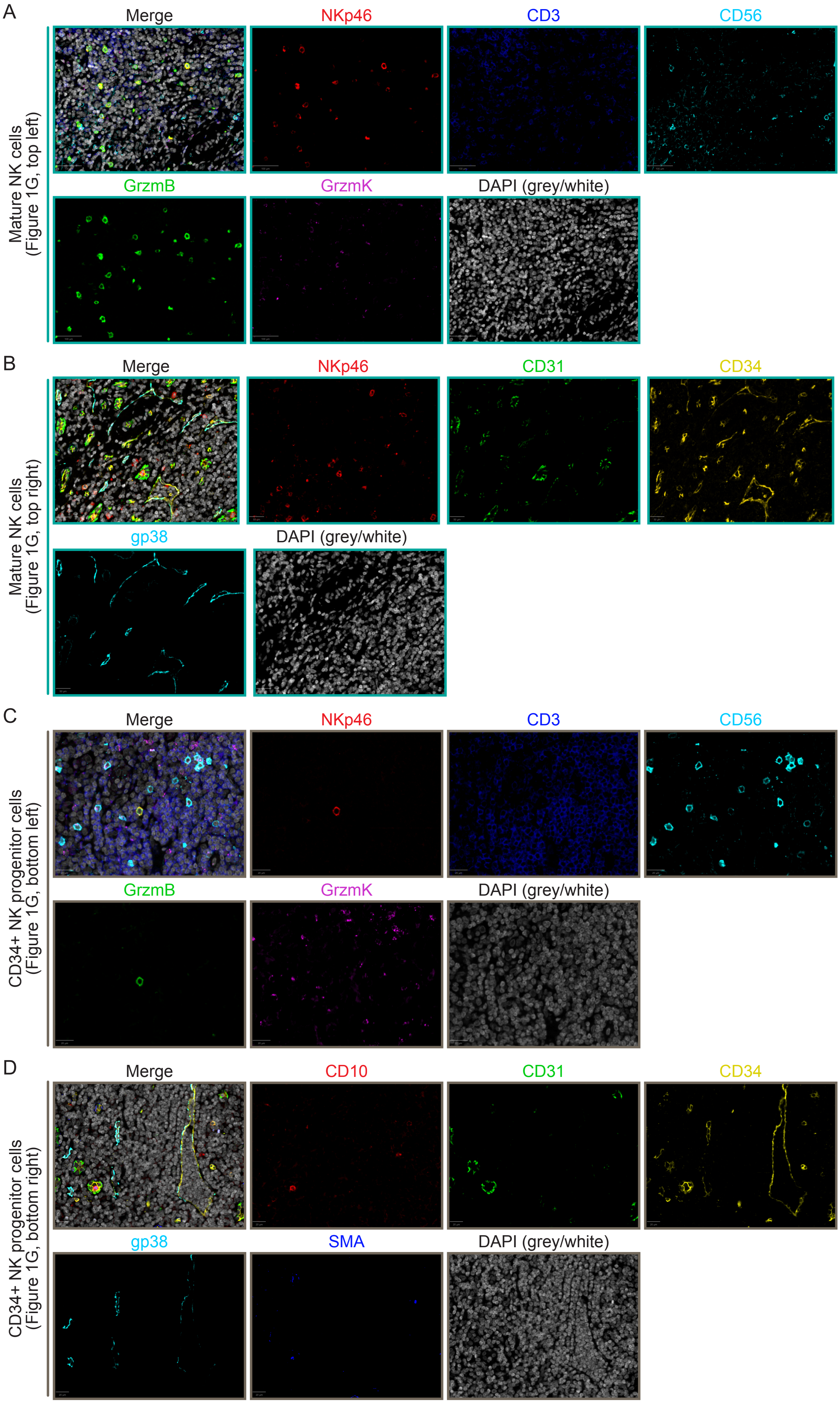

Figure S2: A-D) Merged and single channels of representative CycIF images from Fig. 1G. Images of tonsil NK cell populations were acquired as described in Methods.

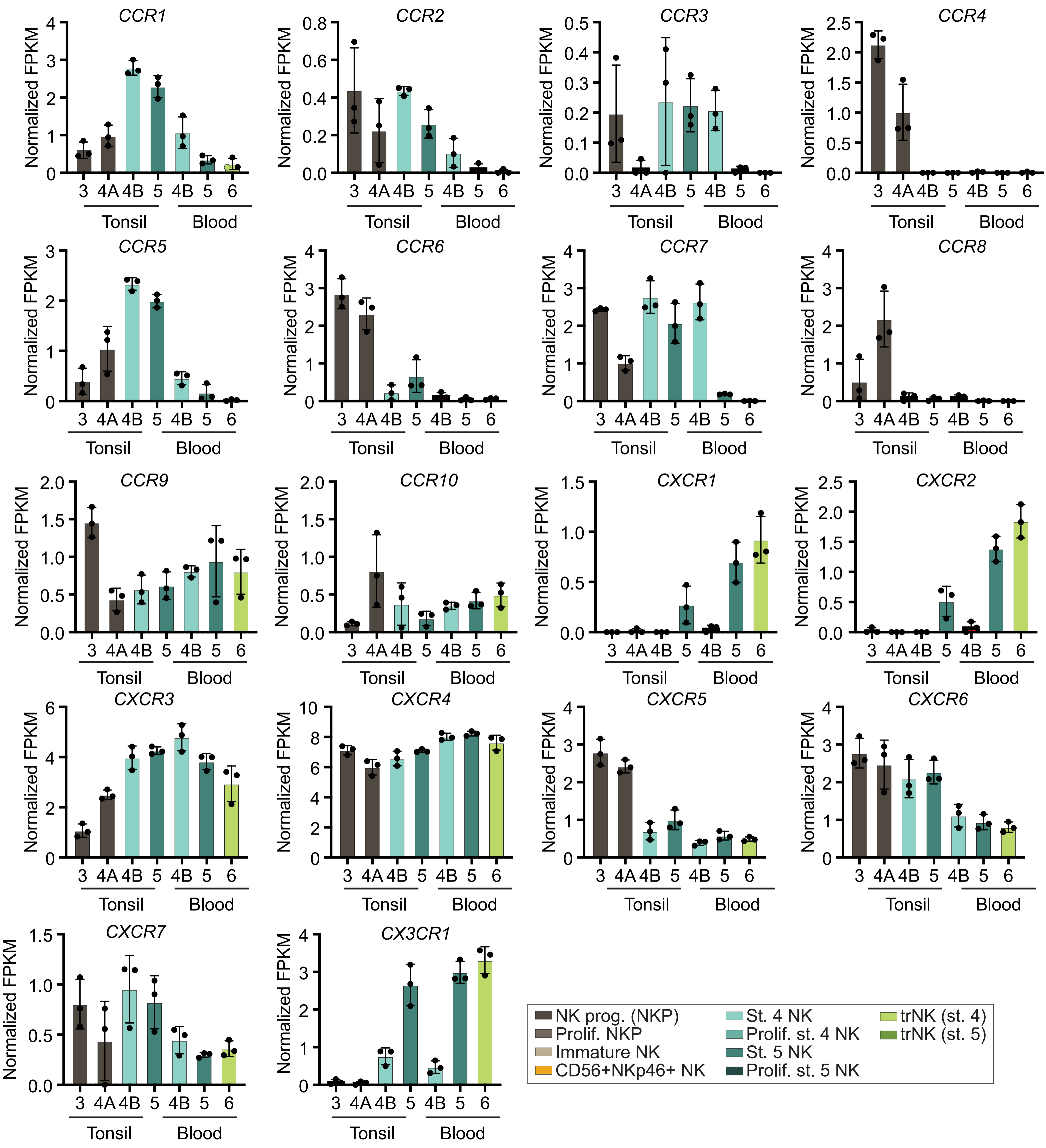

Figure S3: A) Chemokine receptor gene expression (normalized FPKM) of sorted NK cell subsets from peripheral blood and tonsil. NK cell subsets were sorted by FACS and pooled into 3 technical replicates from 12 tonsil and peripheral blood donors for bulk RNA-seq.

Fig. S4 - Solloa et al.

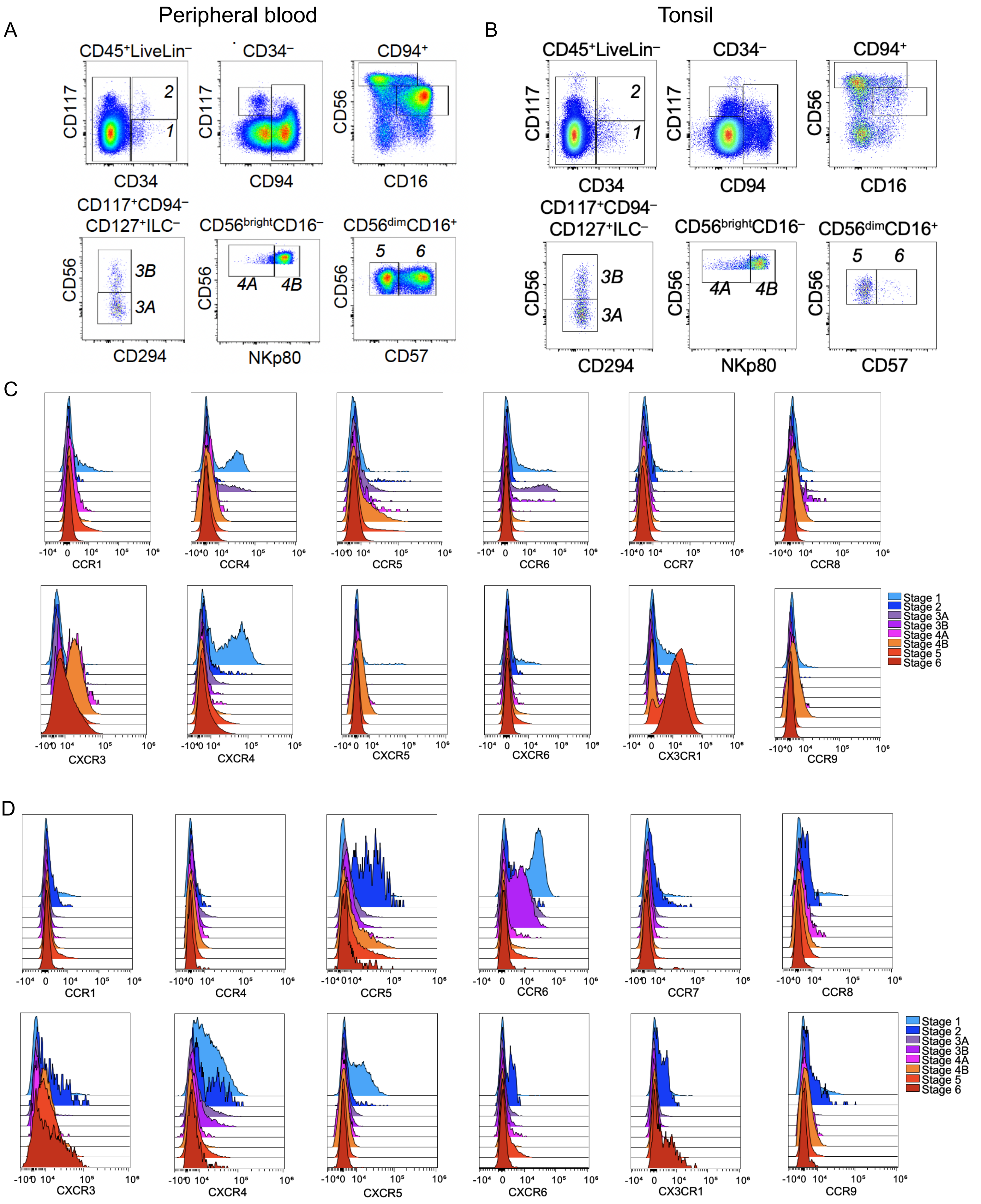

Figure S4: A) Representative flow cytometry gating strategy for measuring peripheral blood and (B) tonsil NK cell subset chemokine receptor expression. C) Tonsil NK cell subset chemokine receptor expression measured by flow cytometry. Each histogram represents 6-7 donors. D) Peripheral blood NK cell subset chemokine receptor expression measured by flow cytometry. Each histogram represents 6-7 donors. Paired peripheral blood and tonsil donors were used for flow cytometric analysis of NK cell chemokine receptor expression.

Table S1

| Round | Marker | Manufacturer | Catalog # | Clone | Removed | Ab Conc. (μg/mL) |
|---|---|---|---|---|---|---|
| **1** | CD56 | RnD | AF2408 | Polyclonal | | 2.5 |
| | CD34 | Novus | NBP2-34713 | QBEnd/10 | | 4 |
| | IL2RB | LSBio | LS-C401487-60 | Polyclonal | | 1.5 |
| **2** | CD49a | RnD | AF5676 | Polyclonal | | 1 |
| | NKp46 | Thermofisher | PA5-79720 | Polyclonal | | 5 |
| | gP38 | Biolegend | 395002 | LpMab-21 | | 5.5 |
| **3** | ICAM1 | RnD Systems | BBA17 | Polyclonal | | 6 |
| | CD20 | BD | 555677 | H1 | | 2.5 |
| | CD127 | Invitrogen | PA5-122138 | Polyclonal | | 5 |
| **4** | CD10 | RnD Systems | AF1182 | Polyclonal | | 6 |
| | CD45 | Invitrogen | 14-9457-82 | CD45-2B11 | | 5 |
| | CD49d | LSBio | LS-B3837-0.05 | Polyclonal | | 7.5 |
| **5** | CD117 | RnD | AF1356 | Polyclonal | Yes | 3 |
| | Ki67 | Fluidigm | 3172024B | B56 | | 5 |
| | CD94 | LSBio | LS-B14034 | Polyclonal | | 10 |
| **6** | CD3 | ABCAM | AB11089 | CD3-12 | Yes | 10 |
| | CXCL12 | RnD Systems | MAB350-SP | 79018 | | 8 |
| | CCL21 | MyBioSource | MBS9411525 | Polyclonal | | 4.8 |
| **7** | CCL3 | RnD Systems | AF-270-NA | Polyclonal | Yes | 10 |
| | CD68 | LSBio | LS-B17033-100 | C68/684 | | 10 |
| | CD3 | ABCAM | AB5690 | Polyclonal | | 1.5 |
| **8** | CX3CL1 | RnD Systems | AF537 | Polyclonal | | 6 |
| | Flt3L | ABCAM | AB52648 | EP1140Y | | 4 |
| | NKp46 | RnD Systems | MAB1850 | 195314 | Yes | 8 |
| **9** | CCL4 | ThermoFisher | MA5-43912 | A4 | Yes | 7.5 |
| | IL7 | Sigma Atlas | HPA019590 | Polyclonal | | 2 |
| | VCAM1 | Novus | NBP2-44615 | VCAM1/843 | Yes | 3 |
| **10** | CD56 | RnD | AF2408 | Polyclonal | Yes | 3 |
| | CCL20 | ThermoFisher | MA5-43912 | A4 | | 10 |
| | GZMK | Invitrogen | PA5-84586 | Polyclonal | | 1.5 |
| **11** | CCL3 | RnD Systems | AF-270-NA | Polyclonal | Yes | 5 |
| | CD34 | Novus | NBP2-34713 | QBEnd/10 | Yes | 5 |
| | GZMB | ABCAM | AB4059 | Polyclonal | | 10 |
| **12** | CX3CL1 | RnD Systems | AF537 | Polyclonal | Yes | 4 |
| | CCL19 | ThermoFisher | TA804261S | OTI2A12 | Yes | 5 |
| | CD94 | ABCLONAL | A6438 | Polyclonal | | 10 |
| **13** | CD117 | RnD | AF1356 | Polyclonal | Yes | 2 |
| | CD122 | LSBio | LS-C401487-60 | Polyclonal | | 3 |
| **14** | ICAM1 | RnD Systems | BBA17 | Polyclonal | | 5 |

| | | | | | | |
|---|---|---|---|---|---|---|
| | VCAM1 | Novus | NBP2-44615 | VCAM1/843 | Yes | 5 |
| | NCR3 | ABCLONAL | A14522 | Polyclonal | | 3.927 |
| **15** | CCL4 | ThermoFisher | MA5-43912 | A4 | Yes | 10 |
| | Pan Keratin | Fluidigm | 3148020D | C11 | | 5 |
| | IL15 | LSBio | LS-B10060 | Polyclonal | | 15 |
| **16** | SMA | Fluidigm | 3141017D | 1A4 | | 5 |
| | Collagen type 1 | Fluidigm | 3169023D | Polyclonal | | 3.75 |
| | CD69 | ABCLONAL | A21174 | Polyclonal | | 15 |
| **17** | CD57 | Biolegend | 359602 | HNK1 | | 15 |
| | CD117 | RnD | AF1356 | Polyclonal | | 4 |
| | CD103 | ABCLONAL | A9934 | Polyclonal | | 14.2 |
| **18** | Ki67 | Fluidigm | 3172024B | B56 | Yes | 6 |
| | CD56 | RnD | AF2408 | Polyclonal | Yes | 5 |
| | CD11c | ABCAM | AB52632 | EPI347Y | Yes | 7 |
| **19** | NKp46 | Innate Pharma | n/a | 8E5B | | 0 |
| | CD11c | Sigma Atlas | HPA004723 | Polyclonal | | 20 |
| | CX3CL1 | RnD Systems | AF537 | Polyclonal | Yes | |
| **20** | CCL19 | ThermoFisher | TA804261S | OTI2A12 | | 2 |
| | CD31 | Novus | NB100-2284 | Polyclonal | | 0.6 |
| | NKp80 | Miltenyi | 130-112-591 | REA845 | | 20 |
| **21** | CD122 | LSBio | LS-C401487-60 | Polyclonal | Yes | 10 |
| **20** | TBET | Biolegend | 644802 | 4B10 | Yes | 10 |
| | CD127 | Invitrogen | PA5-122138 | Polyclonal | Yes | 7 |
| **23** | CD94 | Proteintech | 13332-1-AP | Polyclonal | | 15 |
| | CD8a | Fluidigm | 3162034D | C8/144B | | 5 |

Table S2

| Fluorophore | Marker | Clone/Identifier | Vendor | Catalog # | Dilution |
|---|---|---|---|---|---|
| BUV395 | CD94 | HP-3D9 | BD Biosciences | 743954 | 1:150 |
| BUV496 | CD3 | UCHT1 | BD Biosciences | 612940 | 1:200 |
| BUV496 | CD14 | M5E2 | BD Biosciences | 750381 | 1:200 |
| BUV496 | CD19 | SJ25C1 | BD Biosciences | 612938 | 1:200 |
| BUV563 | CD56 | NCAM16.2 | BD Biosciences | 612928 | 1:100 |
| BUV805 | CD45 | HI30 | BD Biosciences | 612891 | 1:200 |
| BV510 | CD294 | BM16 | Biolegend | 350120 | 1:200 |
| BV650 | NKp44 | p44-8 | BD Biosciences | 744302 | 1:200 |
| BV711 | CD117 | 104D2 | Biolegend | 313230 | 1:100 |
| BV785 | CD103 | Ber-ACT8 | Biolegend | 350230 | 1:150 |
| PE | Ki67 | KI67 | Biolegend | 350504 | 1:100 |
| PE/Dazzle™ 594 | CD34 | 561 | Biolegend | 343534 | 1:150 |
| PE-Cy5 | CD127 | A019D5 | Biolegend | 351324 | 1:150 |
| APC | NKp80 | REA845 | Miltenyi | 130-112-591 | 1:100 |
| AF700 | CD16 | 3G8 | Biolegend | 302026 | 1:150 |
| APC-Cy7 | Zombie NIR | n/a | Biolegend | 423106 | 1:250 |

Table S3

| Panel | Fluorophore | Marker | Clone/Identifier | Vendor | Catalog # | Dilution |
|---|---|---|---|---|---|---|
| all | BUV395 | CD94 | HP-3D9 | BD Biosciences | 743954 | 1:150 |
| all | BUV496 | CD3 | UCHT1 | BD Biosciences | 612940 | 1:200 |
| all | BUV496 | CD14 | M5E2 | BD Biosciences | 750381 | 1:200 |
| all | BUV496 | CD19 | SJ25C1 | BD Biosciences | 612938 | 1:200 |
| all | BUV563 | CD56 | NCAM16.2 | BD Biosciences | 612928 | 1:100 |
| 1 | BV421 | CXCR3 | G025H7 | Biolegend | 353715 | 1:100 |
| 2 | BV421 | CCR5 | J418F1 | Biolegend | 359117 | 1:100 |
| 3 | BV421 | CCR7 | G043H7 | Biolegend | 353208 | 1:100 |
| 4 | BV421 | CXCR4 | 12G5 | Biolegend | 306517 | 1:100 |
| all | BUV805 | CD45 | HI30 | BD Biosciences | 612891 | 1:200 |
| all | BV510 | CD294 | BM16 | Biolegend | 350120 | 1:200 |
| all | BV650 | NKp44 | p44-8 | BD Biosciences | 744302 | 1:200 |
| all | BV711 | CD117 | 104D2 | Biolegend | 313230 | 1:100 |
| all | BV785 | CD103 | Ber-ACT8 | Biolegend | 350230 | 1:150 |
| all | FITC | CD57 | HNK-1 | Biolegend | 359604 | 1:200 |
| 1 | PE | CCR4 | G034E3 | Biolegend | 353418 | 1:100 |
| 2 | PE | CCR9 | L053E8 | Biolegend | 358903 | 1:100 |
| 3 | PE | CCR8 | L263G8 | Biolegend | 360603 | 1:100 |
| 4 | PE | CXCR5 | J252D4 | Biolegend | 356904 | 1:100 |
| all | PE/Dazzle™ 594 | CD34 | 561 | Biolegend | 343534 | 1:150 |
| all | PE-Cy5 | CD127 | A019D5 | Biolegend | 351324 | 1:150 |
| 1 | PE-Cy7 | CCR6 | G034E3 | Biolegend | 353418 | 1:100 |
| 2 | PE-Cy7 | CCR1 | 5F10B29 | Biolegend | 362913 | 1:100 |
| 3 | PE-Cy7 | CX3CR1 | 2A9-1 | Biolegend | 341611 | 1:100 |
| 4 | PE-Cy7 | CXCR6 | K041E5 | Biolegend | 356011 | 1:100 |
| all | APC | NKp80 | REA845 | Miltenyi | 130-112-591 | 1:100 |
| all | AF700 | CD16 | 3G8 | Biolegend | 302026 | 1:150 |
| all | APC-Cy7 | Zombie NIR | n/a | Biolegend | 423106 | 1:250 |

Table S4

| Fluorophore | Marker | Clone/Identifier | Vendor | Catalog # | Dilution |
|---|---|---|---|---|---|
| BUV496 | CD3 | UCHT1 | BD Biosciences | 612940 | 1:200 |
| BUV496 | CD14 | M5E2 | BD Biosciences | 750381 | 1:200 |
| BUV496 | CD19 | SJ25C1 | BD Biosciences | 612938 | 1:200 |
| BUV563 | CD56 | NCAM16.2 | BD Biosciences | 612928 | 1:100 |
| BUV805 | CD45 | HI30 | BD Biosciences | 612891 | 1:200 |
| Pacific Blue | CD31 | WM59 | Biolegend | 303124 | 1:150 |
| BV650 | gp38 (podoplanin) | LpMab-17 | BD Biosciences | 747633 | 1:100 |
| BV711 | CD73 | AD2 | Biolegend | 344026 | 1:100 |
| FITC | CD271 | ME20.4 | Biolegend | 345104 | 1:100 |
| PE | CD56 | 39D5 | Biolegend | 355504 | 1:100 |
| PE/Dazzle™ 594 | CD34 | 561 | Biolegend | 343534 | 1:100 |
| PE-Cy5 | CD106 | STA | Biolegend | 305808 | 1:100 |
| PE-Cy7 | CD90 | 5-E10 | Biolegend | 328124 | 1:100 |
| APC | CD10 | HI10A | Biolegend | 312210 | 1:100 |
| AF700 | CD54 | HA58 | Invitrogen | 56-0549-42 | 1:100 |
| APC-Cy7 | ZombieNIR | n/a | Biolegend | 423106 | 1:250 |

Table S5

| Sample ID | Sex | Age (years) | Inflamation | Pathology Report | Patient notes |
|---|---|---|---|---|---|
| ENT057 | M | 4.75 | yes | acute tonsilitis and lymphoid hyperplasia | |
| ENT059 | M | 4.67 | yes | acute tonsilitis and lymphoid hyperplasia | |
| ENT061 | F | 3.67 | no | lymphoid hyperplasia | |
| ENT066 | M | 3.42 | yes | chronic and acute tonsillitis | |
| ENT067 | M | 4.25 | no | | |
| ENT069 | M | 6.67 | no | | |
| ENT079 | M | 4.9 | no | | |
| ENT088 | M | 6.9 | no | | |
| ENT089 | M | 7.5 | no | | |
| ENT091 | F | 4.5 | yes | | recurring throat infections |
| ENT100 | F | 17 | no | Hypertrophy of tonsils and adenoids | obstructive sleep apnea |
| ENT128 | M | 8.8 | no | Hypertrophy of tonsils and adenoids | recurring throat infections |
